# Atomic resolution ensembles of intrinsically disordered proteins with Alphafold

**DOI:** 10.1101/2025.06.18.660298

**Authors:** Vincent Schnapka, Tatiana I. Morozova, Samiran Sen, Massimiliano Bonomi

**Affiliations:** Institut Pasteur, Université Paris Cité, CNRS UMR 3528, Computational Structural Biology Unit, Paris, France; CNRS, ENS de Lyon, LPENSL, UMR5672, 69342, Lyon cedex 07, France

## Abstract

Intrinsically disordered proteins are ubiquitous in biological systems and play essential roles in a wide range of biological processes and diseases. Despite recent advances in high-resolution structural biology techniques and breakthroughs in deep learning-based protein structure prediction, accurately determining structural ensembles of IDPs at atomic resolution remains a major challenge. Here we introduce bAIes, a Bayesian framework that integrates AlphaFold2 predictions with physico-chemical molecular mechanics force fields to generate accurate atomic-resolution ensembles of IDPs. We show that bAIes produces structural ensembles that match a wide range of high- and low-resolution experimental data across diverse systems, achieving accuracy comparable to atomistic molecular dynamics simulations but at a fraction of their computational cost. Furthermore, bAIes outperforms state-of-the-art IDP models based on coarse-grained potentials as well as deep-learning approaches. Our findings pave the way for integrating structural information from modern deep-learning approaches with molecular simulations, advancing ensemble-based understanding of disordered proteins.

## Introduction

Intrinsically disordered proteins (IDPs) and regions constitute approximately one-third of the human proteome^1^ and play a key role in many biological systems,^2^ yet they are exceptionally challenging to characterize due to their intrinsic dynamic nature. Instead of adopting a single stable structure, IDPs explore a wide continuum of conformations. The structural characterization of these systems is essential for understanding their biological functions and often requires a combination of numerous expensive experimental techniques.^3–6^ For this reason, the ability to predict the conformational behavior of IDPs *in silico* is of paramount importance.

Recently, significant progress toward this objective has been achieved through the development of coarse-grained models for molecular dynamics (MD) simulations.^7–12^ These models enable predicting some of the properties of IDPs with good agreement with experimental data, such as overall chain compaction distribution, and liquid-liquid phase separation propensities. As a result, they provide valuable insight into the global compaction distribution of IDPs in the human proteome.^13^ Furthermore, these remarkable advancements have enabled training deep generative models capable of predicting, from sequence, IDP conformational properties,^12^ intermolecular interactions involving disordered regions^14^ and generating IDPs ensembles that agree with low-resolution experimental data such as small-angle X-ray scattering (SAXS) and Förster resonance energy transfer (FRET), at high throughput.^15–17^

Despite their efficiency, coarse-grained models lack the resolution needed to capture detailed local conformational properties and secondary structure propensities of IDPs, features that are crucial for understanding recognition and binding mechanisms^18^ and interpreting high-resolution experiments such as Nuclear Magnetic Resonance (NMR) spectroscopy. There is also increasing demand for high-throughput generation of atomistic IDP models for drug discovery, where atomic detail is essential.^19–21^ Achieving this level of detail typically requires extensive high-resolution experimental data^3^ as well as all-atom explicit-solvent MD simulations.^22^ However, MD approaches are constrained by high computational cost and limitations in force field accuracy, despite recent advances in IDP-specific force fields^23^ and efficient solvation models.^24,25^ As a result, existing databases of IDP ensembles consistent with experimental data remain sparse, limiting both our understanding of IDP conformational landscapes and the development of accurate deep generative models capable of generating accurate ensembles at atomic-resolution.^26–28^

In parallel, recent advances in deep learning, particularly AlphaFold2,^29,30^ have revolutionized structural biology by predicting highly accurate protein structures from amino acid sequences using data from the Protein Data Bank and co-evolutionary information from multiple sequence alignments (MSAs). Although IDPs lack strong sequence conservation, limiting MSA-based predictions, AF2 has proven valuable in this context.^31^ It can predict disorder,^32^ identify conditional folding via pLDDT scores, ^33^ and provide insights into binding mechanisms.^34–36^ AF2 also outputs residue-residue distance distributions (*distograms*), which measure the local confidence in the AF2 prediction but may also reflect local dynamics and thus aid ensemble modeling. This has sparked growing interest in using AF2 for generating IDP ensembles,^37,38^ including through deep learning models trained on MD simulations.^39,40^ However, the reliability of AF2 for ensemble generation remains debated and requires thorough experimental validation against a wide range of both high- and low-resolution data.

In this work, we leverage the pairwise distograms from AF2 to generate atomistic ensembles of IDPs using bAIes, a Bayesian framework that integrates AF2-derived structural information with prior physico-chemical models. We applied bAIes alongside a computationally efficient random coil model that alone captures key features of disordered proteins. We bench-marked bAIes on several biologically relevant IDPs and evaluated its accuracy against diverse NMR and SAXS data, which probe both local and global conformational properties.^41,42^ Our results show that bAIes efficiently generates accurate ensembles across a spectrum of disorder, from random coils to multi-domain proteins. It is also robust to AF2 prediction errors and can match, or surpass, the accuracy of atomistic molecular dynamics simulations at a fraction of the computational cost. Moreover, bAIes outperforms IDP ensembles generated by state-of-the-art coarse-grained and deep learning-based methods. Overall, bAIes offers a promising and efficient approach for determining accurate atomic-resolution IDP ensembles by leveraging the structural information provided by AF2.

## Results

We assessed the ability of bAIes to model accurate atomistic ensembles of IDPs using a benchmark set composed of 21 proteins (Fig. S1 and Tab. S1). This collection includes IDPs of varying length, amino acid composition, residual secondary structure content, compactness, and overall structural complexity. For most selected systems, NMR chemical shift data are available; in the most favorable cases, this is complemented by an extensive set of NMR observables, including Residual Dipolar Couplings (RDCs), scalar couplings, and Paramagnetic Relaxation Enhancement (PREs) data, as well as SAXS profiles (Tab. S2). This collection of both low- and high-resolution data provide a comprehensive basis for the validation of the ensembles obtained *in silico*.

### An atomic-resolution random coil model for IDPs

To assess the information content provided by AF2, we first developed an efficient method to generate ensembles of IDPs to serve as a structural prior. Specifically, we created a random coil model that excludes any conformational preference or secondary structure propensity. This allows IDPs to theoretically explore the full conformational space of a polymer chain. Similar strategies have been employed to generate initial conformational pools for further refinement using NMR and SAXS data.^43,44^ Our coil model is based on a modified atomistic force field that includes only bonded interactions (bonds, angles, and dihedrals), corrected backbone dihedral distributions, and repulsive Van der Waals terms (Methods; Fig. 1a,b; Fig. S2). Ensemble generation with this model is computationally efficient, requires modest resources, and produces independent structures at a rate comparable to coarse-grained simulations with CALVADOS2^9^ (Fig. S3). This random coil model serves as the structural prior in our bAIes approach and will be integrated with structural information provided by AF2 within our Bayesian framework, as detailed in the next section.

**Figure 1:**
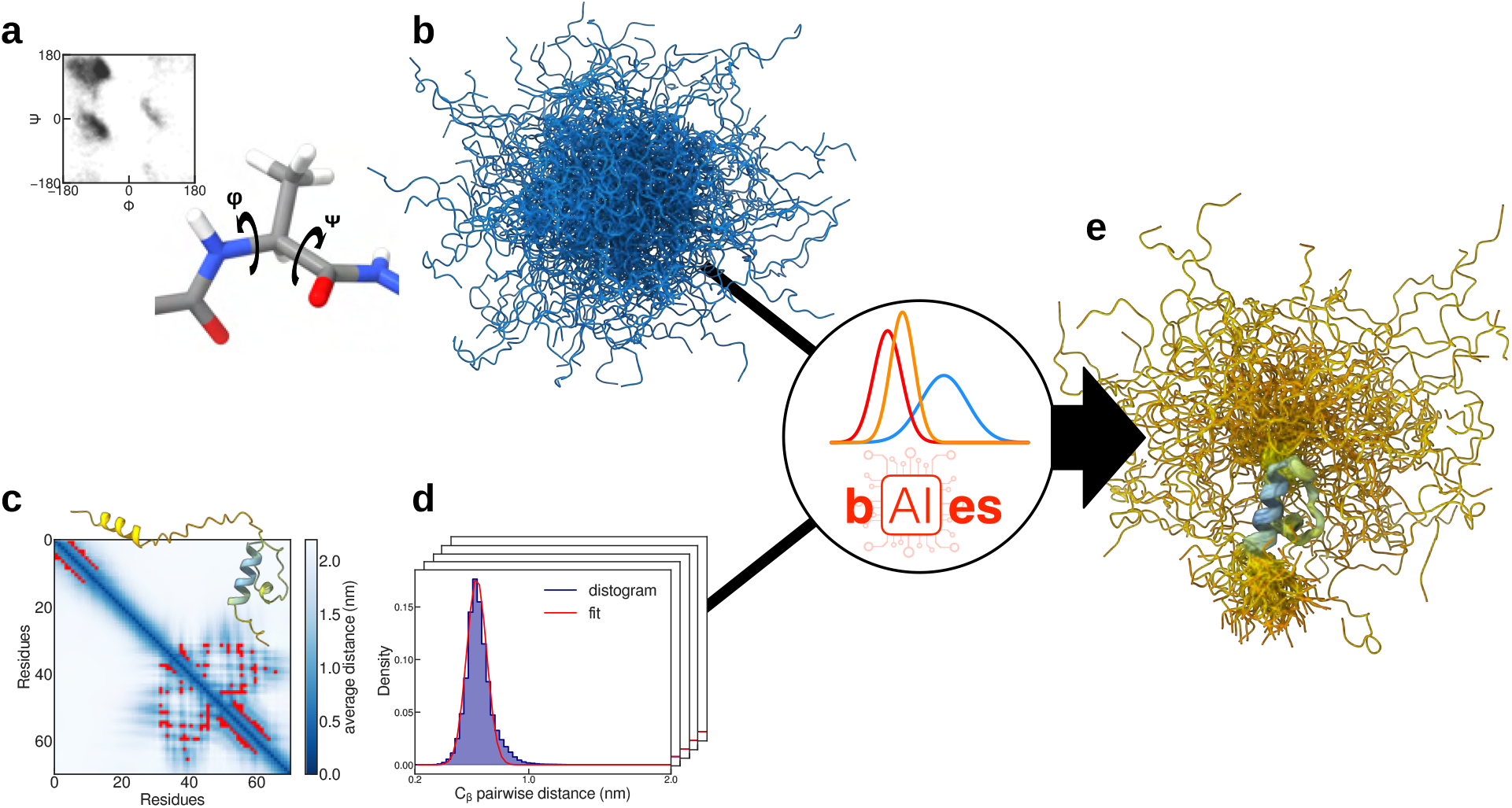
Overview of bAIes. bAIes is a Bayesian modeling framework that combines a random coil model and structural information provided by AF2 to generate accurate ensembles of IDPs at atomic resolution. **a**, The random coil model is a computationally efficient force field with adjusted residue-specific backbone dihedral distributions. **b**, With the coil model only, the system of interest samples the whole conformational space with no structural preference, yielding appropriate prior ensembles. **c**, An Alphafold2 prediction allows the extraction of contact information using distance-based criteria (Methods). **d**, The *C*_*β*_-*C*_*β*_ distograms reporting on contacts are then analyzed and transformed into distance restraints. **e**, These restraints are added on-the-fly to the prior model during the bAIes simulations, yielding ensembles enriched with AF2 information.

### Introducing structural information from AlphaFold

In integrative structural biology, modeling approaches based on Bayesian statistics combine experimental data with prior physico-chemical knowledge through the so-called *data likelihoods* (Methods). These likelihoods ultimately translate into additional structural restraints, which are added to the standard molecular mechanics force fields (the *prior*) to enforce agreement between model and experimental data. Similarly, in bAIes, data likelihoods are modeled from the AF2 distograms, which are distance distributions between *C*_*β*_ atoms (or *C*_*α*_ for glycines). These distributions are provided as a set of equally spaced bins spanning a distance range from 0.2 to 2.2 nm, with the last bin also capturing distances beyond the upper limit. Distograms are used by the AF2 structure module to generate the final model, and as a result, the most probable *C*_*β*_–*C*_*β*_ distance is typically close to the value observed in the resulting structure (Fig. S4).

To integrate AF2 structural information, bAIes likelihood functions were modeled from the AF2 distograms using one Gaussian distribution for each selected pair of residues (Fig. 1c,d). The selected residue pairs are associated with inter-residue contacts (see Methods). To take into account possible errors in the AF2 predictions, we considered the standard deviation of each Gaussian distribution as an unknown (*nuisance*) parameter *σ*_*i*_, with minimum value equal to the width of the corresponding AF2 distogram. A non-informative Jeffreys prior^45^ was used for each *σ*_*i*_ to penalize large errors (Methods). Finally, the AF2 distance-dependent contributions to the force fields derived from the Gaussian likelihoods were added on-the-fly to the prior random-coil model using the open source PLUMED library^46^ (Fig. 1e).

### bAIes captures conformational preferences across the disorder spectrum

For the most disordered proteins with no secondary structure, AF2 is expected to contain only little structural information, as seen in the distogram average distance matrices (Fig. S5). Consequently, very few distance restraints are applied by bAIes to these very dynamic proteins. To demonstrate this point, we used our random coil model and bAIes to generate two ensembles of *Aβ*40 (Fig. 2a,b), a peptide involved in Alzheimer’s disease that is known to exhibit a behavior close to random coil.^47^ To assess the accuracy of our ensembles, we computed the expected NMR chemical shifts, HN-HA ^3^*J* couplings, and NH residual dipolar couplings (Fig. 2c-e). The distribution of these NMR observables along the *Aβ*40 sequence shows that the random coil ensemble already agrees well with available experimental data and, as expected, the incorporation of AF2-derived information through bAIes does not compromise this agreement. This observation was further supported by modeling five other IDPs with no secondary structure, for which no discrepancies were observed between the chemical shifts calculated from the random coil and bAIes ensembles (Fig. 2f and Fig. S6a).

**Figure 2:**
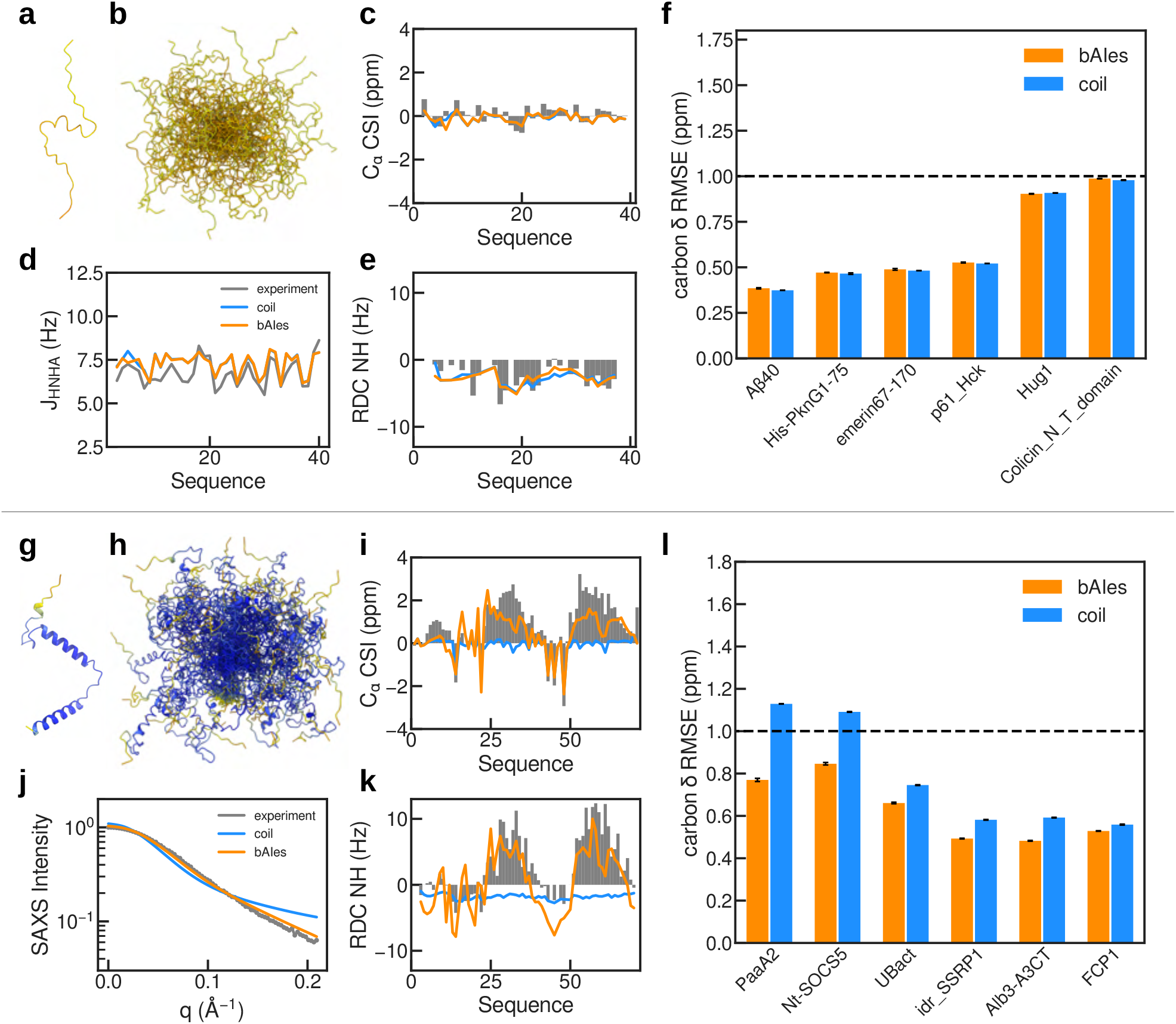
bAIes across the disorder spectrum: from random coil to partially ordered proteins. **a**, Alphafold2 model of A*β*40, colored based on pLDDT. **b**, bAIes ensemble of A*β*40. **c-e**, Validation of the A*β*40 bAIes (orange) and random coil (blue) ensembles with experimental data (grey): **c**, *C*_*α*_ chemical shift index; **d**, ^3^*J*_*HNHA*_ scalar couplings; and **e**, NH residual dipolar couplings. **f**, Average Root Mean Square Error (RMSE) of the random coil (blue) and bAIes (orange) ensembles with respect to experimental backbone carbon chemical shift data (*C*_*α*_, *C*_*β*_, *C*) for six disordered IDPs lacking residual secondary structure. **g**, Alphafold2 model of PaaA2, colored based on pLDDT. **h**, bAIes ensemble of PaaA2. **i-k**, Validation of the PaaA2 bAIes (orange) and random coil (blue) ensembles with experimental data (grey): **i**, *C*_*α*_ chemical shift index; **j**, SAXS profile; and **k**, NH residual dipolar couplings. **l**, Average RMSE of the random coil (blue) and bAIes (orange) ensembles with respect to experimental backbone carbon chemical shift data (*C*_*α*_, *C*_*β*_, *C*) for six partially ordered IDPs. Error bars are obtained from blocking analysis (Methods).

The behavior of disordered proteins and regions cannot always be fully captured by a random coil model. Varying levels of secondary structure propensity often lead to the formation of transient contacts and helical motifs, which are essential to perform specific biological functions. For example, the antitoxin PaaA2, an IDP involved in bacterial stress management, features two helical motifs that mediate toxin binding and are critical for neutralizing its toxic counterpart PaaT, thereby modulating the bacterial stress response.^48^ The AF2 prediction of PaaA2, which accurately identified these motifs (Fig. 2g), is characterized by very high confidence scores (pLDDT *>* 90) even in the highly flexible region between the two helices. The bAIes ensemble correctly captured the flexibility of this protein (Fig. 2h), as demonstrated by the comparison with experimental NMR chemical shifts, RDCs, and SAXS data (Fig. 2i-k). This analysis revealed a clear improvement in the agreement with experimental data with respect to the pure random coil model. In particular, the two helical domains are better represented in the bAIes ensemble, as indicated by larger NMR *C*_*α*_ chemical shifts and positive HN RDC values in the helical domains with respect to the random coil values. In addition, the SAXS profile improved as well with respect to the random coil model, suggesting that additional short-range contacts and better representation of the secondary structure substantially improved the shape distribution of the protein. To confirm that bAIes accurately captures residual secondary structure features, we generated bAIes ensembles for five additional IDPs exhibiting varying levels of structural motifs and for which NMR chemical shift data were available. Compared to the random coil ensemble, we observed a systematic improvement in agreement with the experimental data, which became more pronounced as the amount of residual structural features increased (Fig. 2l and Fig. S6b).

### bAIes is robust to AlphaFold prediction errors

In the previous section, we demonstrated that bAIes can generate accurate ensembles of IDPs across a broad range of disorder. However, AF2 can occasionally mispredict folded elements or secondary structure motifs in IDPs, particularly in regions that fold upon binding. Since AF2 is trained on stable, folded structures deposited in the PDB, the model might be biased toward predicting stable secondary and tertiary structures, even for proteins that are intrinsically disordered in solution. Additionally, because AF2 incorporates evolutionary information from multiple sequence alignments, motifs that fold upon binding to perform specific functions might have a strong sequence conservation signal that can further bias the model toward predicting structured elements, reflecting potential binding-competent states rather than the disordered conformations sampled in solution. To evaluate the robustness of bAIes in such challenging cases, we identified several IDPs for which AF2 models have different levels of errors and we quantified the agreement of our bAIes ensembles against all the available experimental data.

We started by looking at *α*-synuclein, an IDP involved in Parkinson’s disease. Despite the fact that this IDP is very disordered in solution,^49^ AF2 predicted a 90-residue-long helix in the N-terminal domain (Fig. 3a). Surprisingly, the *α*-synuclein bAIes ensemble appeared in reasonable agreement with the available experimental data (Fig. 3b-e). The random coil model alone already captured very well the residue-specific conformational sampling preferences of *α*-synuclein as probed by the NMR chemical shifts and RDCs (Fig. 3c,e). This agreement was only marginally affected by the introduction of AF2 information in bAIes, only a very slight increase in the preference for alpha-helical conformations was observed as seen by the marginal increase in secondary chemical shifts and RDCs in the N-terminal region. Surprisingly, the agreement with SAXS data was improved by adding AF2 data. Taken together, these results indicate that bAIes is robust to errors in the AF2 predictions, thanks to the fact that uncertainties in the AF2 predictions are explicitly modeled using Gaussian likelihoods with variable width (Methods).

**Figure 3:**
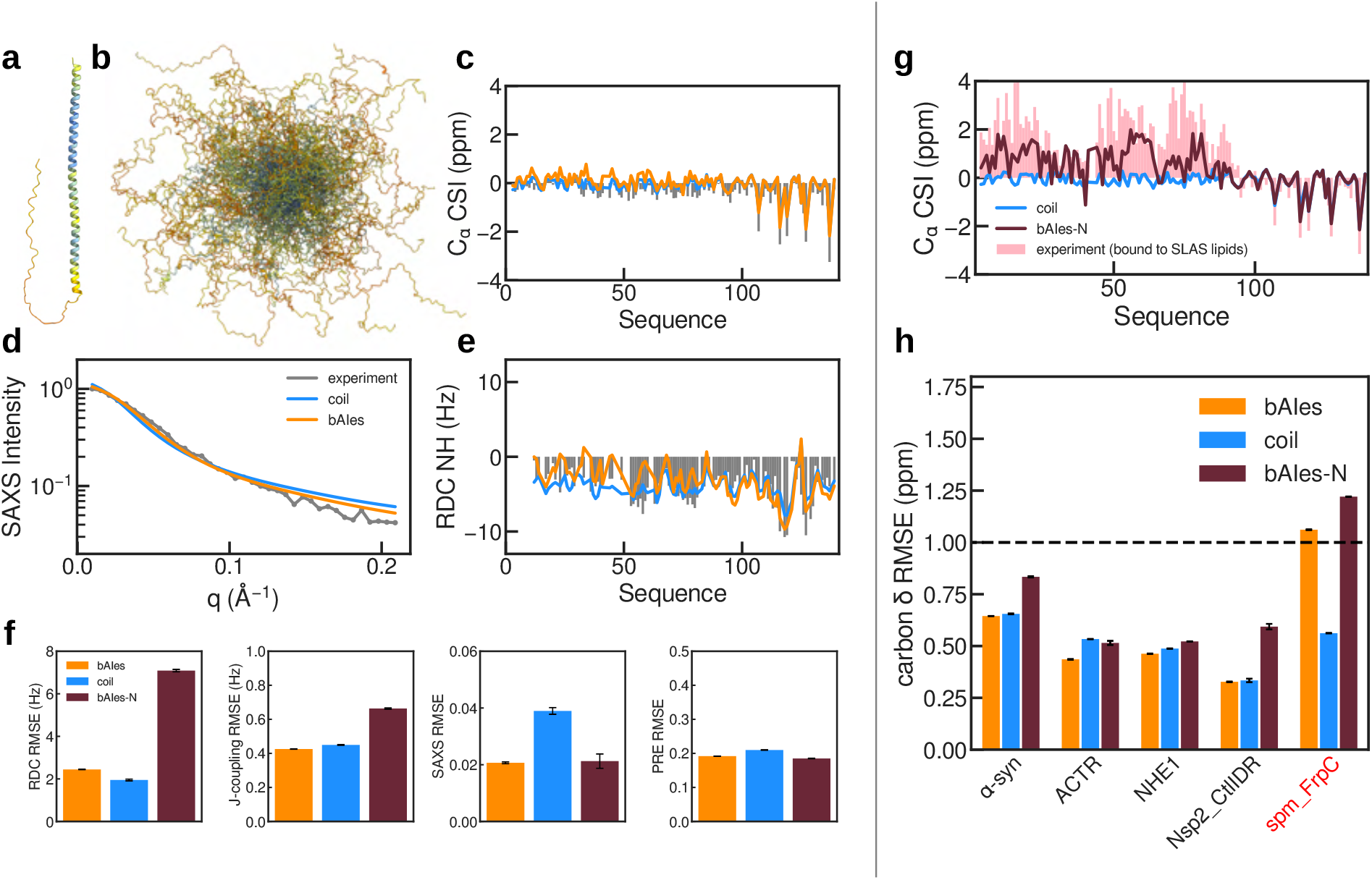
bAIes noise model makes it robust to prediction errors. **a**, AF2 model of *α*-synuclein. **b**, bAIes ensemble of *α*-synuclein. **c-e**, Validation of the *α*-synuclein bAIes (orange) and random coil (blue) ensembles with experimental data (grey): **c**, *C*_*α*_ chemical shift index; **d**, SAXS profile; and **e**, NH residual dipolar couplings. **f**, Average Root Mean Square Error RMSE of the random coil (blue), bAIes (orange), and bAIes-N (dark red) ensembles with respect to RDCs, J-couplings, SAXS, and PREs. **g**, *C*_*α*_ chemical shift index profiles of *α*-synuclein in the random coil (blue) and bAIes-N (dark red) ensembles compared to the experimental chemical shifts (pink) measured on *α*-synuclein in presence of lipids.^103^ **h**, Average RMSE of the random coil (blue), bAIes (orange), and bAIes-N (dark red) ensembles with respect to experimental backbone carbon chemical shift data (*C*_*α*_, *C*_*β*_, *C*) for six IDPs featuring errors in the Alphafold2 predictions. The IDP labeled in red is associated with erroneous predictions of a folded state of the IDP. The error bars are obtained from blocking analysis (Methods).

To test this hypothesis, we designed a modified version of bAIes (bAIes-N), where data likelihoods are derived from a Gaussian fit of the AF2 distograms, without accounting for potential errors in the distribution widths (Methods). The bAIes-N ensembles showed a poorer agreement with the NMR RDCs, J-coupling, and chemical shift data (Fig. 3f,g), in support of our hypothesis. Specifically, the chemical shift profile exhibited higher helical propensities in bAIes-N compared to both bAIes and random coil ensembles. Interestingly, these profiles resemble the experimental conditions in which *α*-synuclein is bound to lipids (Fig. 3g). Despite the poorer agreement with NMR data, the bAIes-N ensemble provided a better fit to the SAXS data compared to the random coil ensemble. This highlights the complementarity of each data type and thus the need for performing a variety of different experiments to validate IDP ensembles. Finally, we did not observe a significant improvement in the agreement with PRE data (Fig. 3f), suggesting that AF2 failed to capture long-range contacts known to exist in this IDP,^50,51^ which are reflected in the PRE profiles (Fig. S7).

To further validate our observations and explore the limitations of our method, we generated ensembles of five additional IDPs known to exhibit conditional folding or to be mispredicted by AF2: ACTR, NHE1, Nsp2 idr, spm FrpC, and the SH3 domain of drk (drkN). The first three IDPs showed a similar trend in the agreement with NMR chemical shift data across random coil, bAIes, and bAIes-N ensembles (Fig. 3h and Fig. S6c), demonstrating that bAIes remains robust while a too high confidence in AF2 prediction (bAIes-N) yields worse agreement against experiments. The case of spm FrpC proved more challenging as AF2 predicts a folded structure, whereas the protein is fully disordered in solution and only folds upon binding calcium ions.^52^ A similar situation arises with drkN, which exists in a dynamic equilibrium between folded and unfolded states in solution, ^53^ yet is also predicted to be folded by AF2. Superimposing the NMR structures of the folded state of drkN and the Calcium-bound folded state of spm FrpC with their respective AF2 models confirmed this observation (Fig. S8). These examples highlight a key limitation of AF2: its inability to account for environmental context or the subtle, transient nature of certain protein conformations. As a result, proteins such as drkN remain particularly challenging.

### bAIes is competitive with state-of-the-art IDPs models

To further put our method to the test, we compared bAIes against state-of-the-art all-atom explicit solvent MD simulations. Specifically, we analyzed large ensembles of A*β*40, PaaA2, *α*-synuclein, and ACTR generated by 30*µ*s-long MD simulations with CHARMM36m,^54^ CHARMM22s,^55^ and amber99SB-disp^23^ force fields, performed with the dedicated *Anton* supercomputer.^56^ Overall, the accuracy of bAIes ensembles was comparable to that of atomistic MD simulations (Fig. 4a). Notably, bAIes outperformed the ensembles generated with the two CHARMM force fields across the entire benchmark set, showing superior agreement with experimental data. When compared to amber99SB-disp, which is widely considered the state-of-the-art force field for IDPs, the performance was generally comparable. Finally, for highly disordered proteins such as A*β*40 and *α*-synuclein, the random coil model already achieved performance comparable to the other IDP models tested, with bAIes offering marginal improvements. This highlights the strength of random coil models in accurately capturing local conformational properties (Fig. 4b-d). A detailed analysis for each protein in our benchmark set is reported in Supplementary Information (Fig. S10).

**Figure 4:**
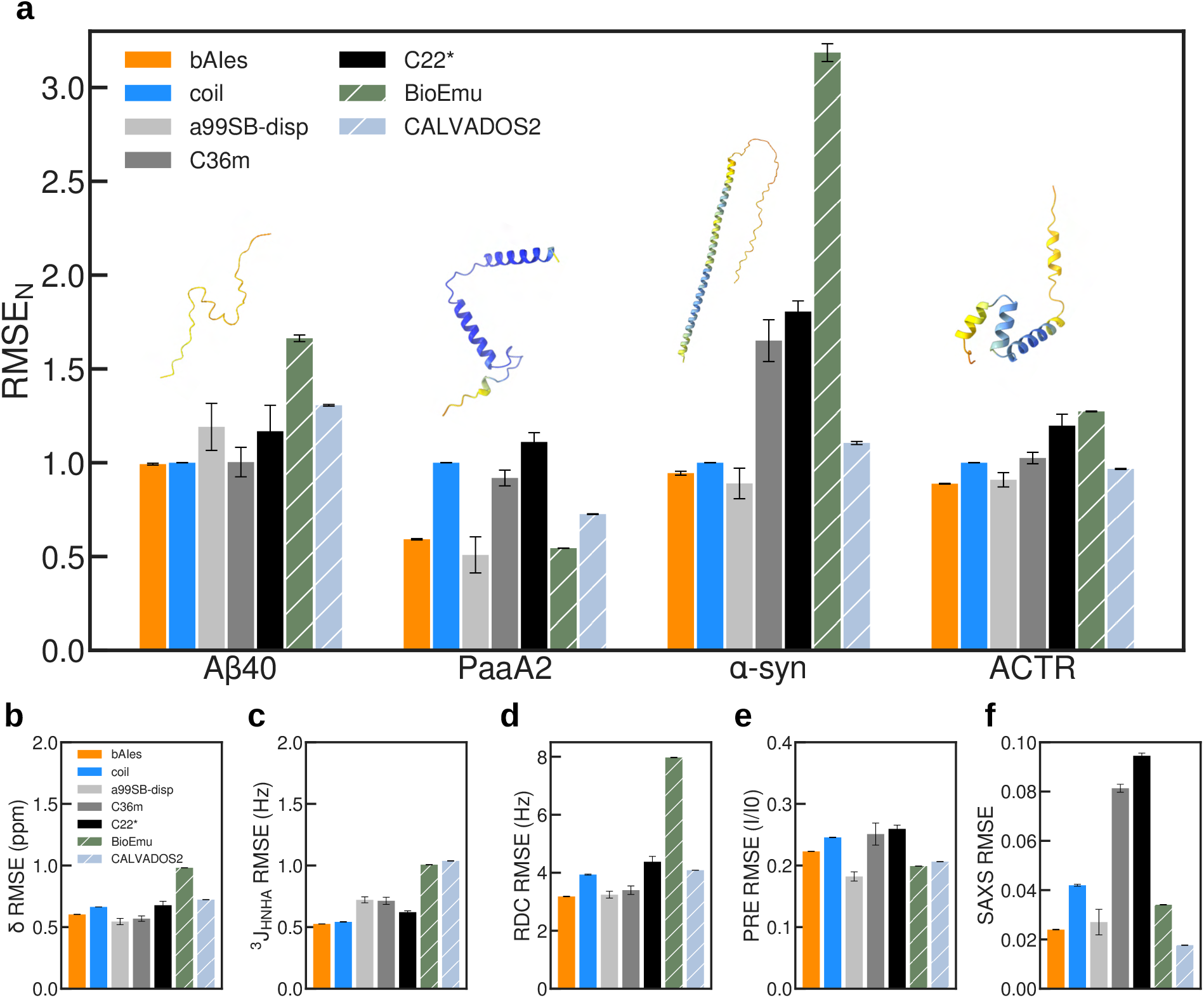
Accuracy of bAIes ensembles against state-of-the-art IDP models. **a**, Normalized Root Mean Square Error (RMSE_*N*_) of the A*β*40, PaaA2, *α*-synuclein, and ACTR random coil (blue), bAIes (orange), amber99SB-disp (grey), CHARMM36m (dark grey), CHARMM22* (black), BioEmu (hatched green) and CALVADOS2 (hatched blue) ensembles. **b-f**, Averaged error across our benchmark set for each data type and force field: **b**, chemical shifts; **c**, J-couplings; **d**, RDCs; **e**, PRE; and **f**, SAXS profile. The error bars are obtained from blocking analysis (Methods).

In addition to classical all-atom MD simulations, we compared bAIes ensembles with those generated using coarse-grained models, represented here by CALVADOS2,^9^ and emerging deep learning-based approaches, exemplified by BioEmu.^39^ These alternative methods offer substantial speed advantages over all-atom simulations. However, bAIes consistently outperformed both in terms of agreement with available experimental data (Fig. 4a). By design, CALVADOS2 is unable to capture local structural motifs or accurately reproduce dihedral angle distributions (Fig. S11). BioEmu, in turn, is limited by the quality of its input embeddings, which are derived from ColabFold predictions. To evaluate this, we compared bAIes and BioEmu ensembles for four IDPs: PaaA2, *α*-synuclein, spm FrpC, and drkN (Fig. S12). While both methods are influenced by the initial AF2 predictions, bAIes, which is explicitly designed to account for prediction errors, yielded ensembles for PaaA2 and *α*-synuclein that are more consistent with experimental data. In contrast, BioEmu tends to produce overly rigid and overconfident ensembles based on its input AF2 structures (Fig. S12a,b), underscoring the strengths of our approach for IDPs. Nonetheless, both BioEmu and bAIes exhibited similar limitations in the cases of drkN and spm FrpC (Fig. S12c,d).

We further analyzed each ensemble by evaluating the agreement with each individual types of experimental data. For chemical shifts, J-couplings, and RDCs, which primarily report on local conformational properties, bAIes performed on par with amber99SB-disp and outperformed all other models (Fig. 4b-d and Fig. S13). This can be attributed to the combination of secondary structure information captured by bAIes and the accurate local conformational sampling of the random coil prior. In terms of SAXS data, both CHARMM force fields were outperformed by bAIes, which achieved a level of agreement comparable to amber99SB-disp. CALVADOS2, which was explicitly optimized using SAXS data, naturally yielded the best agreement with these measurements (Fig. 4f). In contrast, bAIes ensembles were less successful at reproducing PRE data compared to amber99SB-disp (Fig. 4e). This discrepancy likely reflects the limited ability of AF2 to consistently model the transient long-range contacts visible in the PRE profiles of certain IDPs (Fig. S7).

### What contributes to bAIes accuracy?

While we showed in the previous section that bAIes ensembles are in excellent agreement with various types of experimental data, it remains unclear whether this improvement over the coil models arises primarily from AF2-predicted local structural motifs or also involves long-range contacts. To better understand the origin of these improvements, we generated ensembles of A*β*40, *α*-synuclein, ACTR, and PaaA2 using information derived either exclusively from amino acids separated by four residues along the sequence (local contacts) or from long-range predicted contacts. The overall RMSE_*N*_, which quantifies the error across all available chemical shifts, J-couplings, RDCs, PREs, and SAXS data (Methods), indicates that both local and long-range inter-residue contacts contribute to the observed agreement with experimental measurements (Fig. S9). While for A*β*40 all ensembles show comparable agreement with the data, ACTR and PaaA2 clearly demonstrate that incorporating both sources of information yields the most accurate ensembles.

### From bAIes to integrative ensembles of IDPs

We have shown that bAIes can generate structural ensembles that are robust to AF2 prediction errors, achieving an accuracy comparable to state-of-the-art IDP models. We next asked whether this accuracy could be further improved by directly incorporating the experimental data previously used only for validation. To address this question, we applied a Maximum Entropy reweighting technique to refine the ensembles of ACTR, *α*-synuclein, A*β*40, and PaaA2 using extensive datasets of NMR observables and SAXS measurements. This approach minimally modulates the population of a prior ensemble in order to improve the agreement with solution experimental data.

In all systems studied, the reweighted bAIes ensembles outperformed those obtained by refining the CHARMM36m and CHARMM22* ensembles (Fig. 5a). When compared to amber99SB-disp, the reweighted bAIes ensembles were slightly less accurate only in the case of PaaA2. Notably, in the case of *α*-synuclein, where the initial bAIes ensemble was less accurate than amber99SB-disp, reweighting against a large set of experimental data resulted in superior quality compared to the atomistic force field. Furthermore, the bAIes reweighted ensembles still outperformed the reweighted ensembles generated with CALVADOS2 and BioEmu, except for the case of A*β*40, where BioEmu provided the best overall agreement with experimental data across all the IDP models tested.

**Figure 5:**
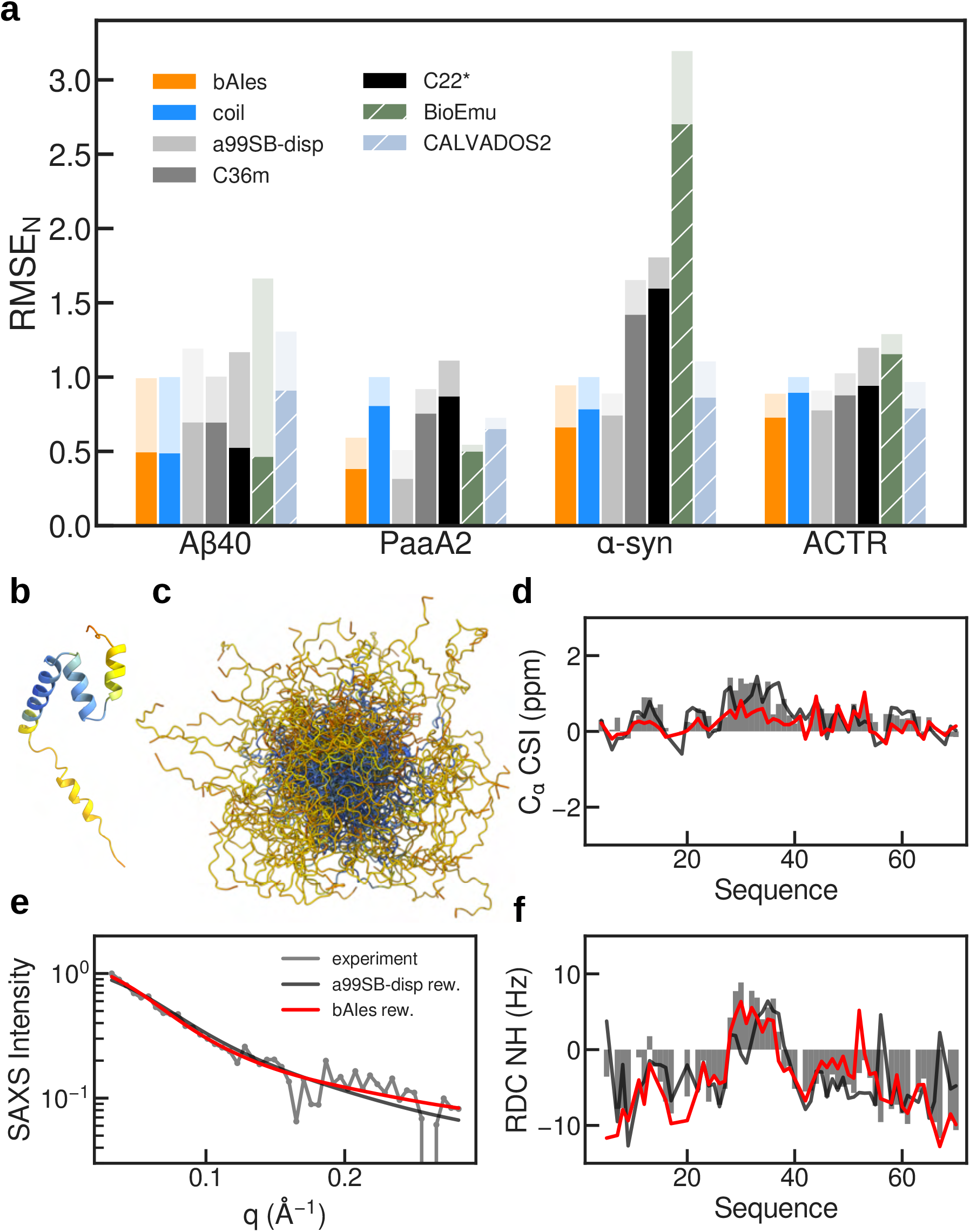
Maximum Entropy reweighting of IDPs ensembles. **a**, Normalized Root Mean Square Error (RMSE_*N*_) of A*β*40, PaaA2, *α*-synuclein, and ACTR for the random coil (blue), bAIes (orange), amber99SB-disp (grey), CHARMM36m (darkgrey), CHARMM22* (black), BioEmu (hatched green) and CALVADOS2 (hatched blue) ensembles for the original (transparent) and reweighted (plain) ensembles. **b**, Alphafold2 model of ACTR, colored based on pLDDT. **c**, bAIes ensemble of ACTR. **d-f**, Validation of the reweighted ACTR bAIes (red) and amber99SB-disp (black) ensembles with experimental data (grey): **c**, *C*_*α*_ chemical shift index; **d**, SAXS profile; and **e**,3N2 H residual dipolar couplings.

Among the IDPs in our benchmark, ACTR was one that contained errors in the prediction of secondary structure, with extra helical propensities that are associated with folding upon binding mechanisms in the function of ACTR (Fig. 5b). Nevertheless, the bAIes ensemble (Fig. 5c) both before and after reweighting shows an overall agreement that competes with amber99SB-disp. Comparing the chemical shift, RDC and SAXS profiles between bAIes and amber99SB-disp confirms this observation, with very similar profiles observed (Fig. 5d-f).

### bAIes ensembles of multi-domain proteins

Finally, we explored the possibility of using bAIes to model multi-domain proteins (MDPs) with folded domains connected by flexible linkers. Since bAIes maintains the integrity of well-defined folded motifs (as seen with drkN and spm FrpC), we expect it to efficiently sample flexible linker conformations while preserving folded motifs. To test this, we generated ensembles of three MDPs: a Green Fluorescent Protein (GFP) chimera composed of two fluorescent proteins separated by a (*GS*)_8_ linker (GS8),^57^ and two polyubiquitin chains, biubiquitin (Ubq2) and triubiquitin (Ubq3).^58^ As expected, in all cases the folded motifs were well predicted by AF2 and preserved in the ensembles generated by bAIes. In addition, our approach allowed efficient sampling of the flexible linkers (Fig. 6a,b,c), showing that bAIes can generate MDPs conformational ensembles without the need to add additional restraints to conserve the folded domains.

**Figure 6:**
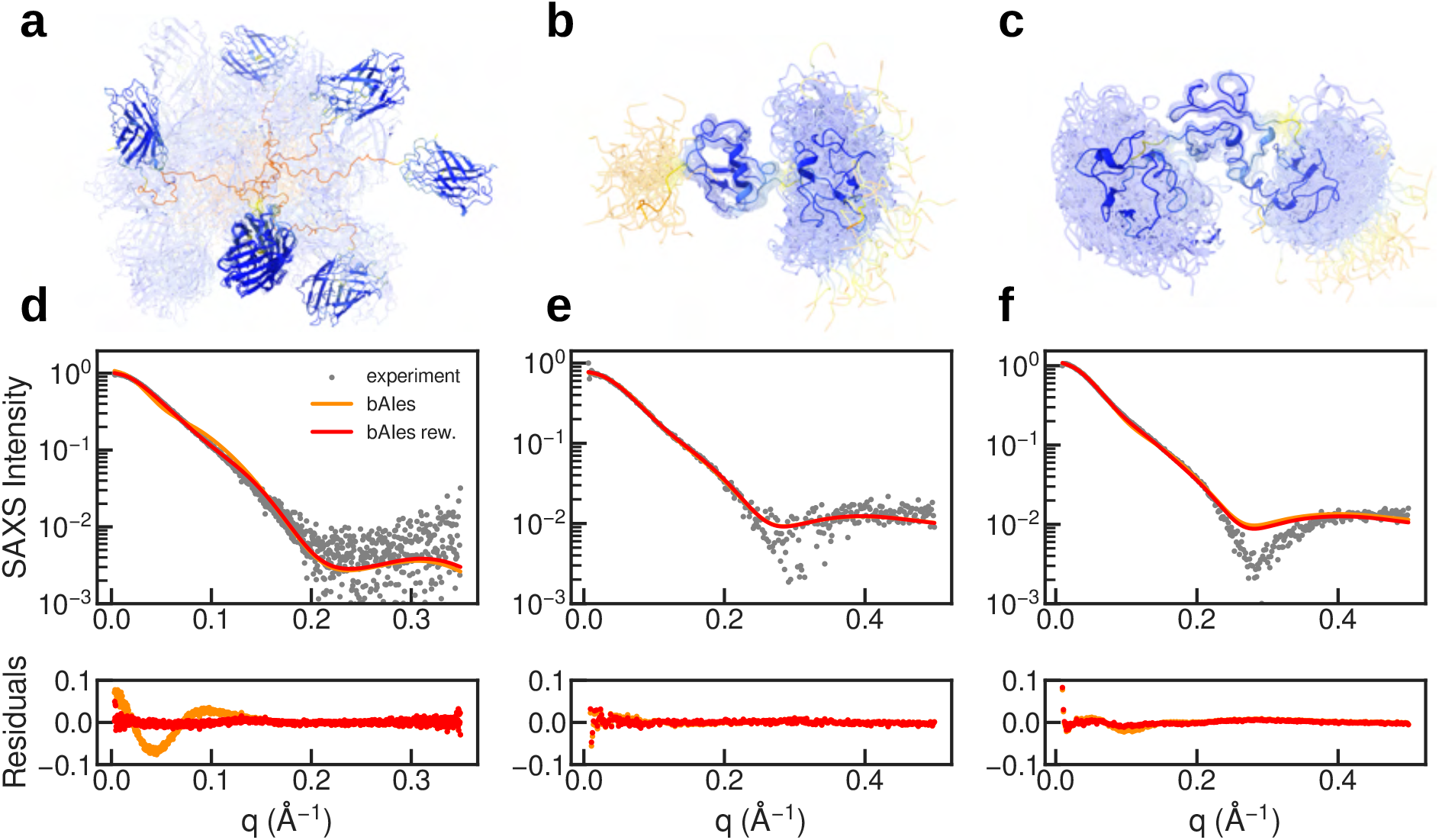
bAIes ensembles of multi-domain proteins. **a, b, c**, bAIes ensemble of GS8, biubiquitin, and triubiquitin, respectively. The models are colored with the pLDDT associated with the AF2 predictions. **d, e, f**, Validation of the bAIes (orange) and reweighted bAIes (red) ensembles using SAXS data (grey) for (**d**) GS8, (**e**) biubiquitin, and (**f**) triubiquitin. Top row: SAXS intensity profiles in logarithmic scale; bottom row: difference between experimental and theoretical profiles (residuals).

We assessed the quality of our ensembles by quantifying the agreement with SAXS data, which provides useful low-resolution information on the overall shape of the protein in solution. The agreement with experimental SAXS profiles, while not perfect, exceeded our expectations given that bAIes only uses AF2 information (Fig. 6d,e,f). In the ensemble of GS8, the two GFP domains sampled a large conformational space, making GS8 relatively extended. Reweighting greatly improved the agreement with SAXS and resulted in more inter-domain contacts (Fig. S14). The polyubiquitins ensembles generated with bAIes showed instead preferred relative domain orientations driven by the inter-domain contacts predicted by AF2. While reweighting slightly improved the agreement with the SAXS data (Fig. S15b,e yellow bars), we wondered whether the predicted inter-domain contacts actually contributed to the match with the experimental profiles. To address this question, we generated bAIes ensembles excluding the inter-domain contacts predicted by AF2. As expected, these ensembles sampled a broader conformational space (Fig. S15a,d): for Ubq2, this had little (negative) impact on the fit with SAXS data (Fig. S15b,c), but for Ubq3, the absence of inter-domain contacts led to poorer agreement (Fig. S15e,f). Furthermore, reweighting ensembles obtained excluding inter-domain contacts failed to reproduce the agreement with SAXS data observed in the original bAIes ensembles (Fig. S15b,e blue bars), suggesting that inter-domain contacts are indeed important for accurately modeling polyubiquitin conformational landscapes.

Although bAIes was not originally designed to model multi-domain proteins, these examples demonstrate its potential for studying this family of proteins. Despite current limitations due the AF2 limited ability to reliably predict transient long-range interactions, our approach holds great promise, especially if enhanced with more accurate physico-chemical priors that can better model inter-domain interactions and integrated with experimental data such as PRE or FRET that probe specific inter-domain contacts.

## Discussions

In this work, we introduced bAIes, a Bayesian framework that leverages the structural information from AF2 to generate accurate atomistic ensembles of intrinsically disordered proteins. Across a range of IDPs with varying residual structure, bAIes matches the accuracy of atomistic molecular dynamics simulations and outperforms coarse-grained and deep-learning-based methods. We attribute this performance to at least three factors. First, the random coil prior captures key IDP properties, as shown by its excellent agreement with experimental data for highly disordered proteins. Second, bAIes effectively incorporates AF2-derived local secondary structure and long-range contact information, improving agreement with NMR and SAXS data over the prior alone. This raises a fundamental question: how does AF2 learn such features despite being trained on folded proteins? One explanation is that it generalizes from semi-ordered loops resolved in PDB structures; more intriguingly, it may have learned general principles of interatomic interactions that also apply to disordered proteins.^59^ Finally, bAIes explicitly models uncertainty in AF2 predictions, preventing overfitting and ensuring robustness to prediction errors.

Leveraging AI-based predictions for ensemble determination is becoming increasingly widespread.^37–39^ In parallel to our work, Metainference^60^ has been applied to integrate AF2-derived structural information with the CALVADOS2 coarse-grained model^9^ for generating IDPs ensembles.^37^

In this approach, the CALVADOS2 force field is minimally adjusted to reproduce the average inter-residue distances derived from the AF2 distograms, generating coarse-grained ensembles that show improved agreement with SAXS data. While conceptually similar to bAIes, this approach differs in several important ways. First, bAIes does not rely on filtering interatomic distances based on AF2 prediction confidence. We highlighted some cases such as PaaA2, where highly flexible regions can exhibit misleadingly high confidence scores (pLDDT *>* 90 and pAE *<* 0.5 nm); applying strong restraints to such flexible regions would artificially rigidify them. Second, bAIes is explicitly designed to capture the dynamic nature of IDPs by leveraging the full distribution of selected AF2 distograms, rather than relying solely on average distance values. Additionally, our method avoids the use of arbitrarily strong structural restraints to enforce secondary and tertiary structure formation, an important advantage when modeling multi-domain proteins. Finally, our method does not require multiple replica simulations and uses a fully atomistic representation that does not require backbone reconstruction. In addition to Alphafold-Metainference, emerging deep generative models also aim to generate protein ensembles from AF2 embeddings. BioEmu,^39^ for instance, is trained on MD data and uses ColabFold^61^ embeddings to emulate backbone ensembles. While promising, we demonstrated that such method is not yet as robust as bAIes in the presence of AF2 prediction errors, and currently yield IDPs ensembles that overall underperform relative to bAIes.

The limitations of our method, which are shared with AlphaFold-Metainference and BioEmu, become evident when AF2 predicts folded states for IDPs. This typically occurs when a protein can adopt multiple states, either under exchange conditions, as in the SH3 domain of drkN, or in distinct environments, such as spm FrpC. These mispredictions are due to the fact that AF2 was not designed to account for the effect of the environment on protein folding. Addressing this limitation would require either training structure prediction models on proteins characterized under standardized conditions or developing models that explicitly incorporate environmental parameters, both solutions currently hindered by the limited availability of experimental data. In addition, AF2 is inherently limited in its ability to capture the transient nature of certain intramolecular contacts, which are often critical and better captured by experimental techniques such as PRE and FRET.^62**?**^

A potential route for improving our model is to incorporate additional energy terms into the random coil model to account for structural features not captured by AF2. One such strategy could involve parameterizing residue-specific potentials, similar to those implemented in modern IDPs coarse-grained models.^9,11^ Additionally, integrating bAIes with enhanced-sampling techniques, such as metadynamics,^63^ could facilitate the exploration of regions of conformational space that are rarely sampled in standard simulations. For example, in the case of drkN, bAIes simulations do not sample the unfolded states, due to the fact that AF2 predict a folded conformation. By using metadynamics to bias the sampling toward less populated, unfolded conformations, one can enhance the exploration of the conformational space. These enhanced ensembles can then be refined using reweighting approaches guided by experimental data to recover the correct equilibrium between folded and unfolded states. Another idea to tackle the existence of distinct conformational states is to exploit multiple distograms generated by running AF2 with modified multiple sequence alignments (MSAs).^64,65^ One could envision a future extension of bAIes that incorporates multiple distograms, derived from different MSAs, to better capture sparsely populated folded states within the conformational landscape of IDPs. Looking ahead, the growing number of structure prediction tools like Alphafold3^30^ and Boltz2^66^ could further expand the range of applications of our approach. For instance, bAIes could be applied to model dynamic RNA molecules, to efficiently sample small-molecule conformations within protein binding pockets, or to investigate the effects of post-translational modifications on IDPs structural properties.

Generating accurate atomic-resolution ensembles of IDPs is becoming increasingly important. These proteins play essential roles in cellular signaling, regulation, and molecular recognition, functions that depend critically on their structural flexibility and ability to engage in dynamic interactions with multiple partners. Atomic-level insights are thus essential for elucidating the molecular mechanisms underlying these behaviors. In this context, bAIes offers a valuable and computationally efficient approach for generating accurate, atomic-resolution IDP ensembles. These ensembles not only advance fundamental research but also support a wide range of applications. First, they can provide preliminary structural descriptions of novel systems, guiding the design of future experimental studies. Second, they serve as large conformational pools that can be validated against, or refined using, diverse experimental datasets. Atomistic detail is also crucial for small-molecule docking, where scoring functions commonly employed in drug discovery rely heavily on atomic-level accuracy.^21^ Finally, there is a growing need for ensembles that are consistent with both high- and low-resolution experimental data to serve as *ground truth* for training machine learning models capable of predicting structural ensembles directly from sequence.^67^ In summary, bAIes provides a computationally efficient approach for generating accurate atomistic ensembles of IDPs. The method is implemented in the open-source PLUMED library (www.plumed.org),^46^ making it readily accessible to the broader computational science community.

## Methods

### General Alphafold settings

Unless stated otherwise, Alphafold^29^ 2.3.2 was run on a local cluster with default settings. MSAs were performed on Uniref90^68^ and MGnify^69^ and template search was performed on the pdb70^70^ with HHsearch^70^ and HHblits.^71^ MSAs on BFD^29^ and Uniref30^72^ were performed with jackhmmer.^73^ To maintain maximum accuracy and applicability for multidomain proteins, we systematically used the AF2 prediction with the highest global pLDDT score among the five generated models.

### Development of the random coil model

The amber99SB-ILDN force field^74^ was modified to create a computationally efficient random coil model. We first defined a simplified force field *E*_FF_ composed by all bonded terms as well as the repulsive part of the Van der Waals potential. In order to do so, these interactions were switched off at a distance cut-off set to 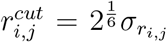 for each pair of atoms *i* and *j*. This distance cutoff corresponds to the energy minimum of the Lennard-Jones potential:

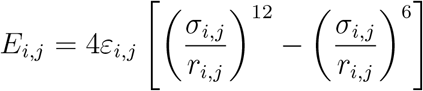

where *r*_*i,j*_ in the distance between atoms *i* and *j*; 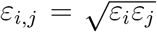 and *σ*_*i,j*_ = 0.5(*σ*_*i*_ + *σ*_*j*_) are the Lennard-Jones parameters. An offset was added to *E*_*i,j*_ to preserve the continuity of the energy function at the cutoff distance. The backbone dihedral distributions *P*_FF_(*ϕ*_*i*_, *ψ*_*i*_) obtained with *E*_FF_ were then corrected to account for the absence of non-bonded interactions. To do so, we computed for each residue type *i* a correction MAP (CMAP) term,^75^ defined as:

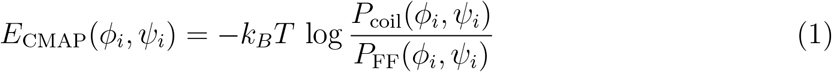

where *P*_coil_(*ϕ*_*i*_, *ψ*_*i*_) is the target residue-specific random coil dihedral distribution, *T* the temperature of the system, and *k*_*B*_ the Boltzmann constant.

To obtain an estimate of the residue-specific random coil distributions *P*_coil_(*ϕ*_*i*_, *ψ*_*i*_), we used a well-established approach that consists in extracting dihedral distributions corresponding to residues in coil regions from structures deposited in the PDB database.^43,44,76,77^ In our study, we selected a pool of protein X-ray structures deposited in the PDB database before January 1st 2024. These structures were filtered to an RFree parameter below 0.25 and sequence identity below 50%, resulting in a total of approximately 30,000 structures. The structures were then analyzed with DSSP^78^ and residues belonging to alpha-helical and beta strand secondary structures were discarded to generate residue-specific (*ϕ, ψ*) distributions representative of random coil residues. Pre-proline residues were analyzed separately,^43^ resulting in a total of 40 *P*_coil_(*ϕ*_*i*_, *ψ*_*i*_) distributions, which were stored on a 24×24 grid following the format adopted by the CHARMM force fields.^75^

To compute *P*_FF_(*ϕ*_*i*_, *ψ*_*i*_), we used well-tempered metadynamics^79^ on 40 tripeptides with amino acid sequence equal to AXA or AXP, where X is one of the 20 canonical amino acids. While the pre-proline residues were considered separately, we arbitrarily chose Alanine as the neighboring residue. Since in the simplified force field *E*_FF_ the non-bonded interactions are turned off, we assumed that this set of tripeptides give a sufficiently accurate estimate of the residue-specific backbone dihedral angles distributions of the central aminoacid, which was later confirmed by the good performance of our random coil model. For each tripeptide sequence, the metadynamics collective variables used were the backbone dihedral angles Φ and Ψ. Gaussians with height equal to 1.2 kJ/mol and widths of 0.2 radians for both dihedrals were added every 1 ps using a bias factor of 10. Metadynamics simulations were carried out with LAMMPS 2 Aug. 2023^80^ equipped with PLUMED 2.8^46^ for 100 nanoseconds each. Finally, *E*_CMAP_(*ϕ*_*i*_, *ψ*_*i*_) were calculated from Eq. 1, stored on a grid, and added to the simplified force field *E*_FF_ to obtain our final random coil model:

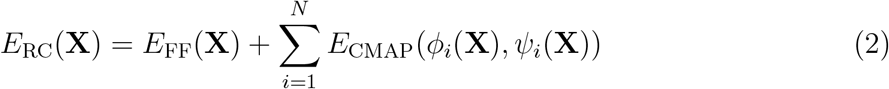

where **X** are the coordinates of the system, and *N* the number of residues.

### bAIes theory

Our bAIes approach is based on a Bayesian inference framework that estimates the probability of a model *M*, defined in terms of its structure **X** and other parameters, given the information available about the system, including prior physico-chemical knowledge and newly acquired data.^81^ The posterior probability *p*(*M* | *D*) of model *M* given data *D* and prior knowledge is expressed as:

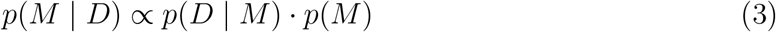

where the likelihood function *p*(*D* | *M*) represents the probability of observing data *D* given *M*. This function quantifies the agreement between the observed data and the model and accounts for all sources of uncertainty and error. The term *p*(*M*) denotes the prior probability of model *M* based on existing knowledge. In the following, we specify each component of this general Bayesian framework as applied in our bAIes approach.

#### The data

In integrative structural biology applications, *D* typically represents a collection of experimental data points, which may originate from different experimental techniques, such as NMR, SAXS, or cryo-electron microscopy.^81–83^ In bAIes, *D* represents the structural information provided by AF2, defined as a collection of distances *{d*_*i*_*}* between *C*_*β*_ atoms (or *C*_*α*_ in the case of glycines). Each data point *d*_*i*_ represents the average distance computed from the corresponding AF2 distogram. Two selection criteria were applied: (i) distances between residues separated in sequence by less than 3 aminoacids were discarded; (ii) pairs with most probable distance in the AF2 distogram greater than a residue-pair specific cutoff^84^ were discarded to retain only potential inter-residue interactions.

#### Data likelihood

For each pair of selected residues, a Gaussian likelihood was used:

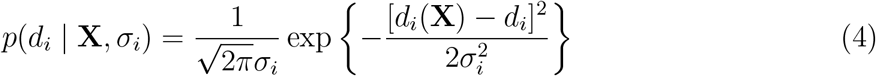

where *d*_*i*_(**X**) is the distance between a pair of *C*_*β*_ atoms (or *C*_*α*_ in the case of glycines) in structure **X** and *σ*_*i*_ an error parameter that modulates the deviation between observed and computed data. Under the assumption of independent data points, the total likelihood for a set of of *M* points *D* = *{d*_*i*_*}* can be factorized as:

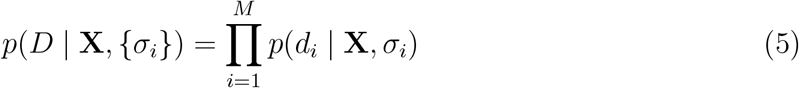

#### Priors

In bAIes, the error parameters *σ*_*i*_ are treated as unknown variables and are determined by sampling the posterior distribution. To penalize large errors, we used an informative Jeffreys prior:

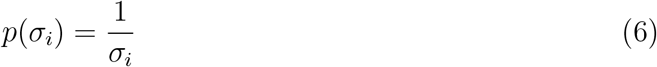

Additionally, the width of the corresponding distogram, calculated by fitting this distribution with a Gaussian function, was used as the minimum value 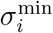 for each uncertainty parameter. This choice allows us to handle cases where AF2 predictions are inaccurate in the same way as we would treat outlier experimental data points in other Bayesian inference approaches for structural biology.^83^ The random coil model previously described was used as structural prior:

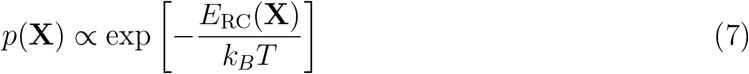

#### Marginalization

To avoid sampling the parameters *σ*_*i*_, we marginalized the corresponding likelihood distributions along with their uncertainty priors. For each data point, the resulting marginal data likelihood is:

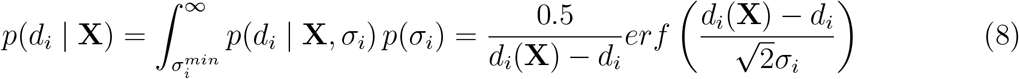

#### Simplified noise model

To assess the effect of our noise model in presence of AF2 prediction errors, we defined a simplified version of bAIes (bAIes-N) in which the *σ*_*i*_ parameters are modeled by a delta Dirac function 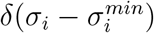, which results in the following marginal data likelihood:

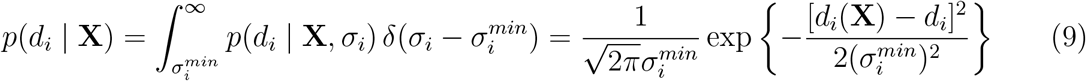

#### The bAIes hybrid energy function

After defining all the component of our approach and marginalizing the uncertainty parameters, we obtain the final bAIes posterior distribution:

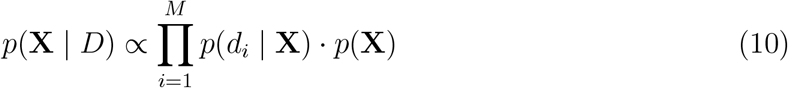

where the marginal data likelihood is given by Eq. 9 and the structural prior by Eq. 7. To sample the posterior, we define the associated bAIes hybrid energy function as:

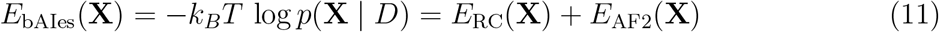

which is composed by our random coil model *E*_RC_ and the spatial restraints *E*_AF2_ that incorporate the structural information provided by the AF2 distograms. Finally, models are sampled by Molecular Dynamics under the bAIes hybrid energy function defined in Eq. 11.

### Details of the random coil model and bAIes simulations

The simulations were carried out using LAMMPS^80^ version 2 Aug. 2023 and the development version of PLUMED.^46^ The setup, minimization, and production steps described below are common to random coil model and bAIes simulations.

- For each protein, the LAMMPS topology and input files are generated from GROMACS inputs prepared in vacuo with the top ranked AF2 model and using the amber99SB-ILDN force field.^74^ The conversion from LAMMPS to GROMACS is performed using InterMol 0.1.0 and ParmEd 4.2.2.^85^ The amber99SB-ILDN force field is then transformed into our random coil model by a custom Python script, which generates the final LAMMPS input and topology files.
- Prior to production, an energy minimization is performed using the Polak-Ribiere version of the conjugate gradient algorithm (LAMMPS default). Stopping tolerances of 10^*−*4^ and 10^*−*6^ *kcal*.*mol*^*−*1^.*A*^*−*1^ are used for energy and force respectively.
- Production simulations are carried out using the Bussi-Donadio-Parrinello thermostat^86^ set at a temperature of 298.1 K, a damp parameter of 100 fs and a time step of 1 fs. The coordinates are saved every 100 ps, producing semi-independent conformations due to the rapid motion of the protein in this framework (Fig. S3). The final ensembles contain 20000 structures for the 18 studied IDPs and 10000 structures for the 3 multi-domain proteins (GS8, Ubq2 and Ubq3).
- For bAIes simulations, AF2 predictions are first performed. Then, a preprocessing python script reads the AF2 distograms and generates the necessary PLUMED files. The bAIes forces added by PLUMED are updated every two timesteps.

### CALVADOS2 and BioEmu ensembles

CALVADOS2 simulations were carried out using openMM 8.2.0^87^ following the procedure described in Tesei *et al*.^9^ using a Langevin integrator at a temperature of 298 K, a friction coefficient of 0.01 *ps*^*−*1^ and a time step of 5 fs. The pH was set to 7.4 and the beads coordinates were saved every 100 ps. A total of 20,000 conformations were generated for each protein. The all-atom model was then reconstructed with Pulchra^88^ and protons were added with GROMACS 2022.5.^89^

BioEmu ensembles were generated using the protocol described in Lewis *et al*.,^39^ aiming for 30,000 conformations with default settings. The resulting conformations were filtered to remove non-physical structures using the built-in BioEmu option. Finally, side chains and protons were added with hpacker^90^ and GROMACS 2022.5,^89^ respectively.

### NMR and SAXS data back-calculation

Chemical shifts were predicted with SPARTA+.^91^ The ^3^*J*_*HNHA*_ couplings were calculated with a Karplus relationship reported by Vogeli and Bax,^92^ with Karplus parameters of 7.97, -1.26 and 0.63 Hz respectively. The ^3^*J*_*CC*_ couplings were calculated with a Karplus relationship reported by Li *et al*.^93^ with Karplus parameters of 1.61, -0.93 and 0.55 Hz respectively. The PREs were calculated using a rotameric ensemble of MTSL paramagnetic side chains,^94^ assuming a correlation time of 4 ns for the electron-nucleus interaction vector. The PREs were converted to intensity ratios assuming a transverse relaxation rate of 4 s^*−*1^ and an INEPT time of 10 ms. The RDCs were calculated with PALES^95^ using a local alignment window of 15 residues. A local alignment window implicitly neglects the contributions of long-range interactions to the calculated RDC, but allows a rapid convergence of the calculated data.^96^ For each ensemble, the alignment scaling factor for the calculated RDC was optimized on the calculated and experimental data points 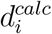 and 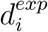 that verify 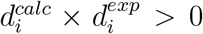. This allows us to optimize agreement with the experimental data while conserving physically sound scaling factors in cases where the experimental profile strongly deviates from the calculations.

The SAXS profiles were calculated with Pepsi-SAXS^97^ using the following procedure. We first computed, for each ensemble conformer, a theoretical profile using a fixed scale factor of 1 and no offset. For the 18 IDPs, the values of the hydration shell contrast *dρ* and effective atomic radius *r*_0_ were set for all conformers at 3.34 *·* 10^*−*3^ *e*Å^*−*3^ and 1.68 Å, respectively, as reported in a previous optimization procedure for IDPs.^98^ For GS8 and polyubiquitins, we scanned for each conformer 31 values of *dρ* between 3.34 *·* 10^*−*5^ and 3.34 *·* 10^*−*2^ *e*Å^*−*3^, and 11 values of *r*_0_ between 1.45 and 1.77 Å. For each choice of parameters, an ensemble-averaged profile was then computed across all conformers along with an optimal scale factor and offset to minimize the *χ*^2^ with respect to the experimental SAXS profile. Finally, the *dρ* and *r*_0_ pair that resulted in the minimum value of *χ*^2^ across all the parameters tested was selected.^99^

### Maximum Entropy reweighting

To improve agreement with ensemble-averaged experimental data in solution, we refined our *in silico* structural ensembles using the Maximum Entropy approach developed by Cesari *et al*.^100^ Briefly, this refinement approach aims at minimally modifying the weights of each conformation in a prior ensemble to improve the fit with the observed data. This approach requires optimizing one or more regularization parameters that reflect the confidence in the experimental data. We used a protocol recently proposed^22^ to seamlessly determine both individual regularization parameters, once for each experimental data type, as well as a global one. The protocol is based on reaching a target effective reduction of the initial ensemble, or Kish ratio, equal to 0.1. Experimental data used for reweighting include all the available chemical shifts, J-couplings, PREs, RDCs and SAXS profiles reported in this work. The optimal RDC scaling factors as well as the SAXS scaling and background parameters were determined by an iterative process of reweighting and parameter optimization.^98^

### Details of ensemble validation

The quality of all ensembles generated in this study was assessed based on their agreement with the available experimental data. Five types of experimental data were used in this work: chemical shifts, J-couplings, RDCs, PREs, SAXS profiles. For chemical shifts, J-couplings and PREs, different experimental datasets are available corresponding to different nuclei, nuclei pair and paramagnetic label location in the sequence, respectively. The available data for each protein are reported in Tab. S2.

For each experiment type and dataset composed of *N* data points, the Root Mean Square Error (RMSE) between the experimentally measured values 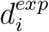 and the ensemble-averaged values 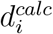 computed from the ensemble generated with the force field *ff* was defined as:

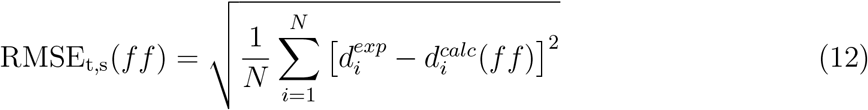

where *t* is the type of experiment (chemical shift, J-coupling, RDC, PRE, or SAXS), and *s* is the dataset associated with the type of experiment, such as *C*_*α*_ chemical shift or *H*_*N*_ -*H*_*α*_ J-coupling.

The RMSE for a given type of experiment was defined as the average RMSE across multiple datasets:

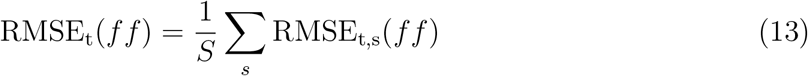

where *S* is the number of datasets associated with a given experiment type. Finally, to provide a global assessment of a given force field *ff*, we defined a global normalized Root Mean Square Error RMSE_*N*_ (*ff*) relative to the random coil reference *rc*, as follows:

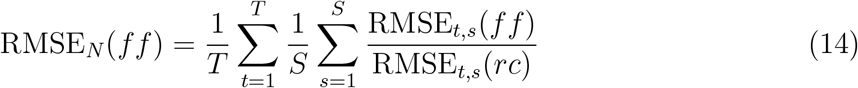

where *T* is the total number of experimental data types considered. The normalization against the random coil model provides a unitless metric that allows us to provide an overall evaluation of the quality of an ensemble using all the types of experimental data available.

Error bars on the various RMSE metrics introduced above were obtained using blocking analysis. Each trajectory was first divided into five non overlapping blocks, and all analysis were repeated separately on these five blocks. The error bar associated with a given RMSE corresponds to the standard error of the mean across the five blocks.

## Software and data availability

bAIes is implemented as a part of the PLUMED-ISDB module^101^ in the GitHub master branch of PLUMED (https://github.com/plumed/plumed2).^46^ The LAMMPS topologies and PLUMED input files used in our benchmark are available in PLUMED-NEST (www.plumed-nest.org), the public repository of the PLUMED consortium,^102^ as plumID:25.014. Scripts to prepare bAIes simulations as well as a complete tutorial are available at https://github.com/COSBlab/bAIes-IDP.

## Acknowledgments

The authors would like to thank Paul Robustelli and Guillaume Bouvignies for useful discussions. This project has received funding from the European Research Council (ERC) under the European Union’s Horizon 2020 research and innovation programme (Grant agreement No. 101086685 – bAIes).

