## Supplementary material for "Atomic resolution ensembles of intrinsically disordered proteins with Alphafold": Supp Info

#### Supporting Information

Vincent Schnapka,[1] Tatiana I. Morozova,[1,2] Samiran Sen,[1] and Massimiliano Bonomi[*,1]

[1]*Institut Pasteur, Université Paris Cité, CNRS UMR 3528, Computational Structural Biology Unit, Paris, France*

[2]*Current address: CNRS, ENS de Lyon, LPENSL, UMR5672, 69342, Lyon cedex 07, France*

*

### Supplementary Text

#### Effect of AlphaFold2 parameters on IDPs predictions

To better understand how to run our predictions and which information is relevant and should be taken for bAIes, we first evaluated whether different choices of the AF2 setting parameters could have an impact on the determination of the distograms and on the final AF2 structural models. Our analysis showed that the use of structural templates and a moderate reduction of the MSA database had little effect on the structural predictions of IDPs (Fig. S16). On the other hand, keeping default MSA settings and using structural templates ensures good accuracy for folded domains.[1] We therefore chose to use the deepest MSA along with structural templates in our analyses to ensure maximum accuracy for proteins containing folded domains.

#### Selection of AlphaFold2 distograms

We then assessed whether all distance distributions are equally relevant for constructing our bAIes approach. To this end, we performed simulations of two disordered proteins: A$\beta$40 and $\alpha$-synuclein using both our random coil model and CALVADOS2, and compared the resulting distance distributions with those obtained from all-atom explicit solvent MD simulations using the amber99SB-disp force field,[2] as well as with the AF2 distograms. For A$\beta$40, the amber99SB-disp MD simulations showed a stronger correlation with the AF2 distograms compared to the random coil and CALVADOS2 models (Fig. S17a). Notably, the classical all-atom simulations identified contacts within A$\beta$40 that were also predicted by AF2 but missed by the other models (Fig. S17b). In contrast, for $\alpha$-synuclein, AF2 wrongly predicts a helix in the N-terminus region, yielding poor correlations in this region between AF2 distances and simulations (Fig. S17c). On the other hand, for the C-terminal region of $\alpha$-synuclein, the average $C_\beta$-$C_\beta$ pairwise distances generated by the three models correlated equally well with those derived from the AF2 distograms (Fig. S17d). This analysis indicates that not all the AF2 distograms are necessarily relevant, unless they exhibit information on residual structural propensity or inter-residue contacts, and even this information can be erroneous as shown with $\alpha$-synuclein. From this, we concluded that the relevant residue pairs should be selected with a strict contact-based criterion and that the derived forces should account for possible errors in the AF2 prediction.

#### Relation between distograms and predicted alignment error

Finally, we analyzed the variability in the widths of the AF2 distograms and observed a correlation between the distance standard deviation and the predicted alignment error (pAE). This finding suggests that the spread of the distograms is, to some extent, linked to the uncertainty in determining the relative position of two residues (Fig. S4b). As expected, highly dynamic disordered regions and loops displayed high pAE values, reflecting the inherent uncertainty in the relative positions of residues within these flexible segments. These observations suggest that there might be, to some extent, a correlation between the predicted position uncertainties and the dynamic nature of some residues. This motivated us to combine the AF2 distograms with our random coil model to improve the accuracy of the generated structural ensembles.

### Supplementary Figures

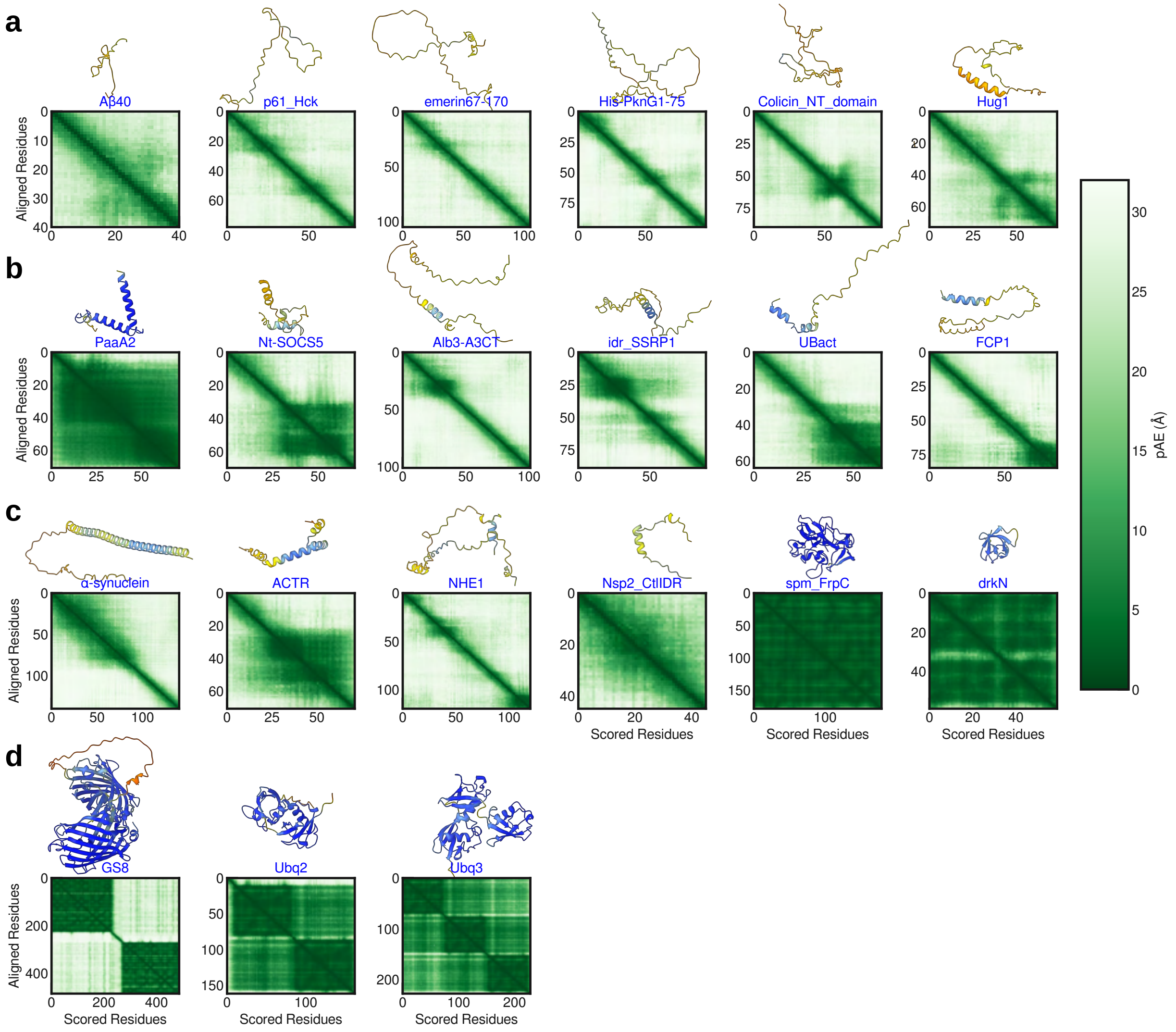


**Supplementary Figure 1: Confidence in AF2 predictions across our entire benchmark set.** **a-d**, AF2 Structural models and predicted alignment error (pAE) matrices for (**a**) fully disordered IDPs, (**b**) IDPs with residual secondary structure, (**c**) IDPs with prediction errors, and (**d**) multidomain proteins. The models are colored based on the pLDDT score: blue (high confident, pLDDT> 90), light blue (confident, 90 > pLDDT > 70), yellow (low confident, 70 > pLDDT > 50), and orange (very low confident, pLDDT < 50).

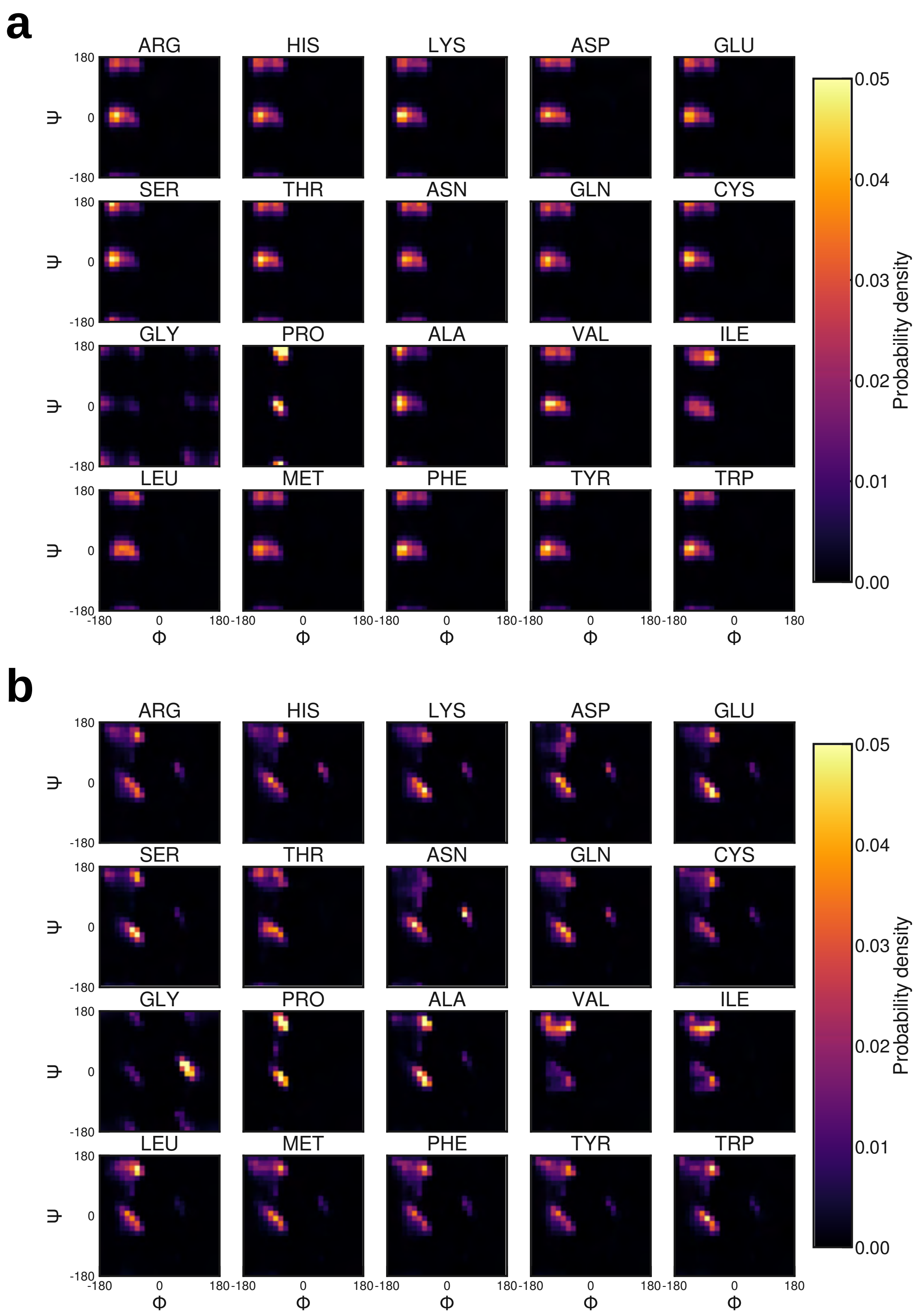


**Supplementary Figure 2: Dihedral distributions of the 20 aminoacids with and without CMAP corrections. a,b**, Ramachandran plots of the 20 aminoacids using our coil model (**a**) without and (**b**) with CMAP corrections. The distributions were obtained using well-tempered metadynamics on AXA tripeptides (Methods).

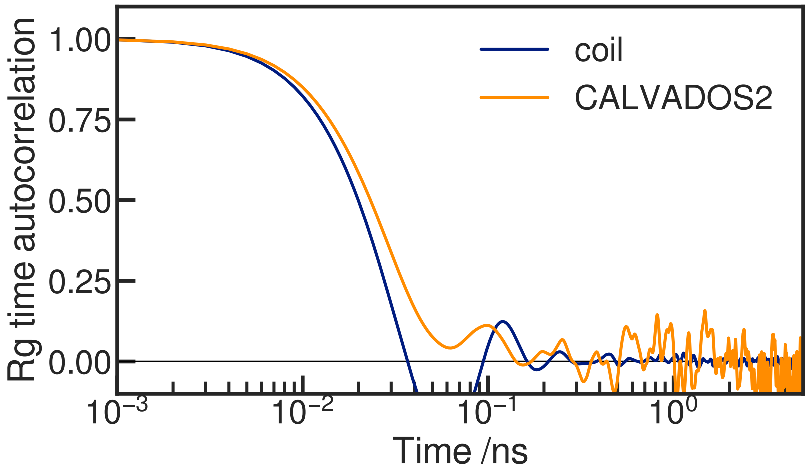


**Supplementary Figure 3: CALVADOS2 and random coil model sampling efficiency**. Time autocorrelation function of the radius of gyration for $A\beta40$ simulations performed with CALVADOS2 (orange) and our random coil model (blue).

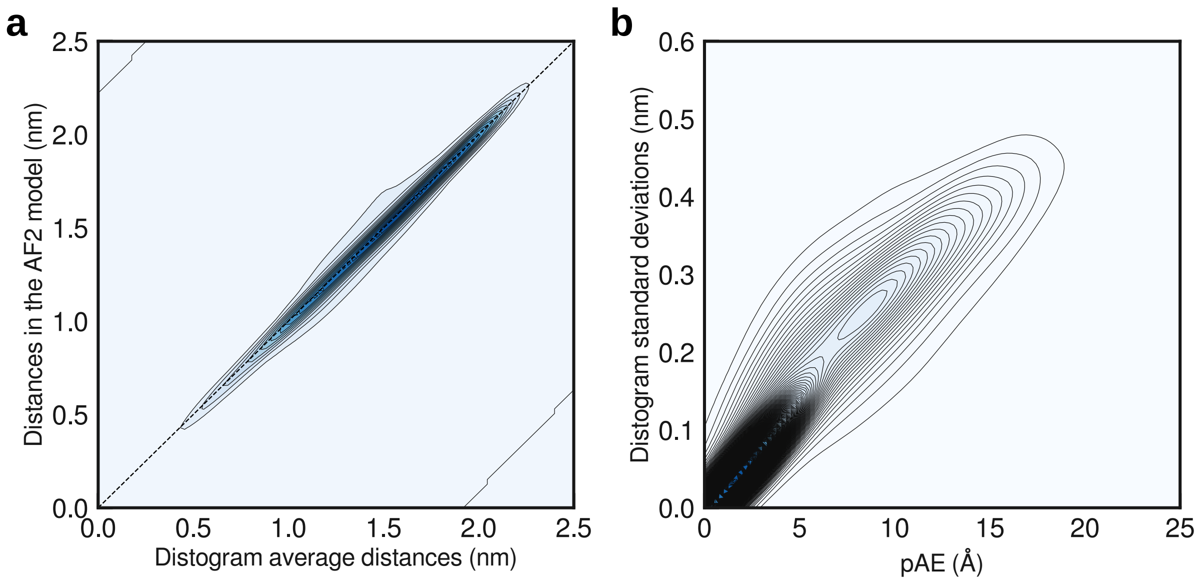


**Supplementary Figure 4: Analysis of AF2 distograms**. **a**, Kernel density estimation of the distribution of measured distances in the AF2 top pLDDT model against the average distance in the corresponding distogram. **b**, Kernel density estimation of the distribution of distogram standard deviations against the associated pAE. Only distograms with negligible information in the last bin (<10%) and corresponding to amino acids pairs separated in sequence by at least 4 residues were considered. These distributions were computed over the entire benchmark set of IDPs used in this work.

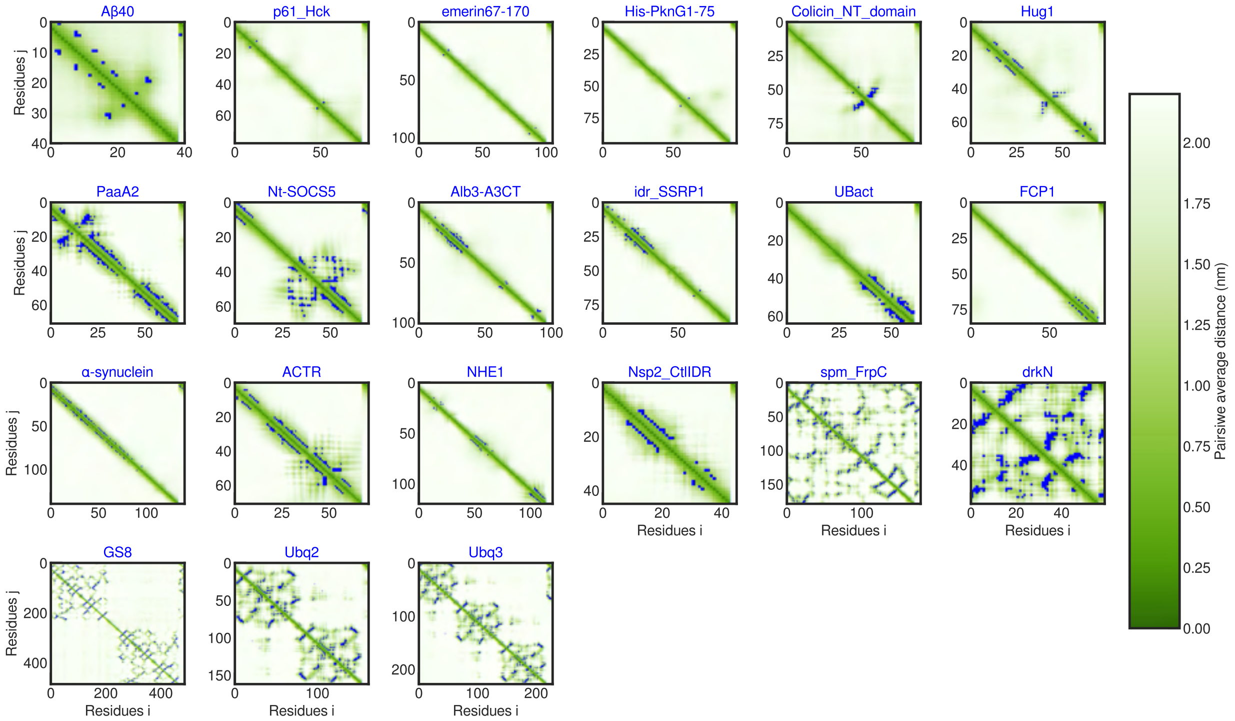


**Supplementary Figure 5: Distogram analysis and contact selection**. Average distances matrix calculated from the AF2 distograms with red squares indicating the residue pairs used in bAIes, based on our contact selection criterion (Methods).

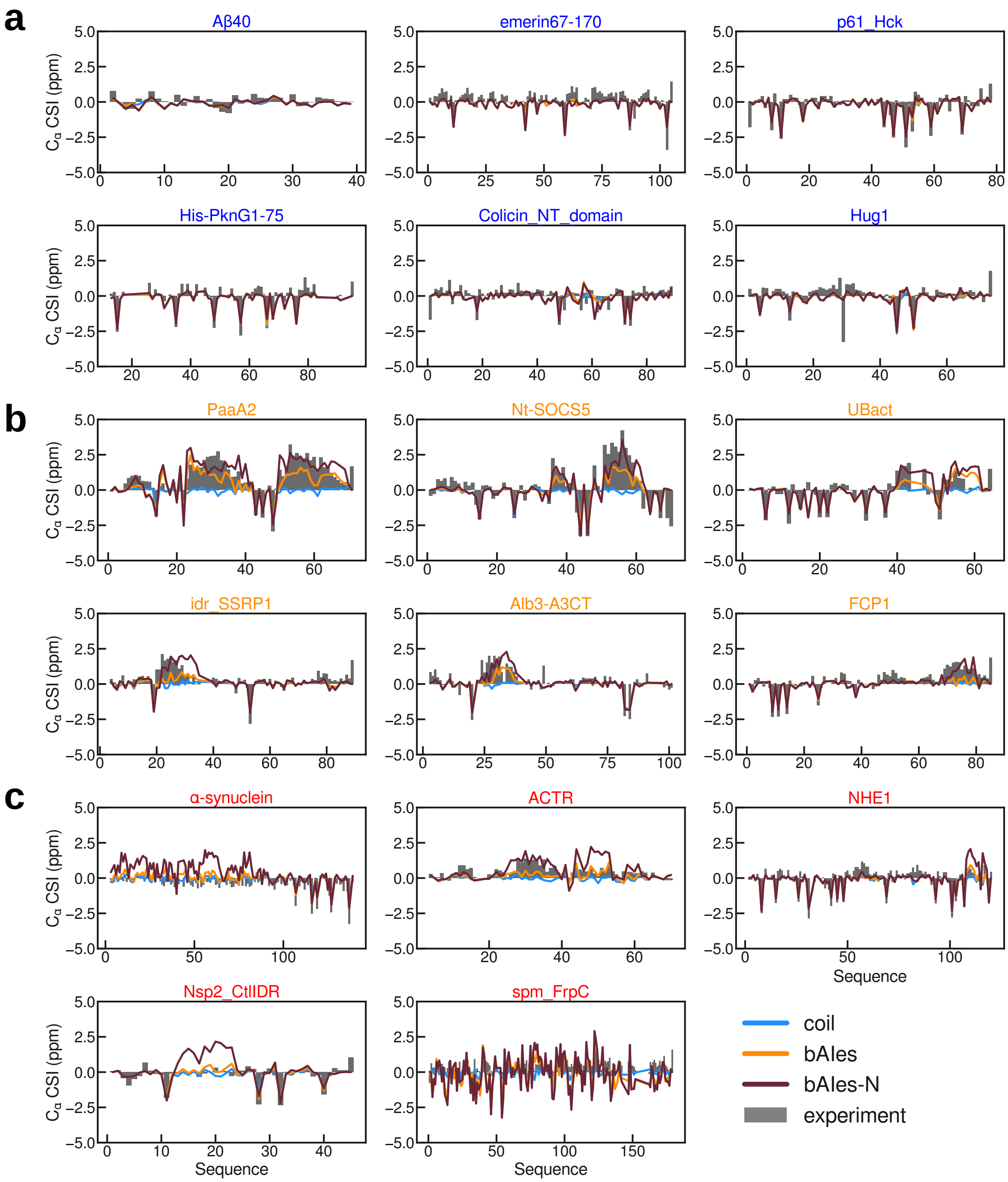


**Supplementary Figure 6: Analysis of backbone $C_\alpha$ chemical shift index (CSI).** (**a**) Fully disordered IDPs, (**b**) IDPs with residual secondary structure, and (**c**) IDPs with prediction errors. Experimental CSI in grey, random coil, bAIes, and bAIes-N in blue, orange, and dark red, respectively.

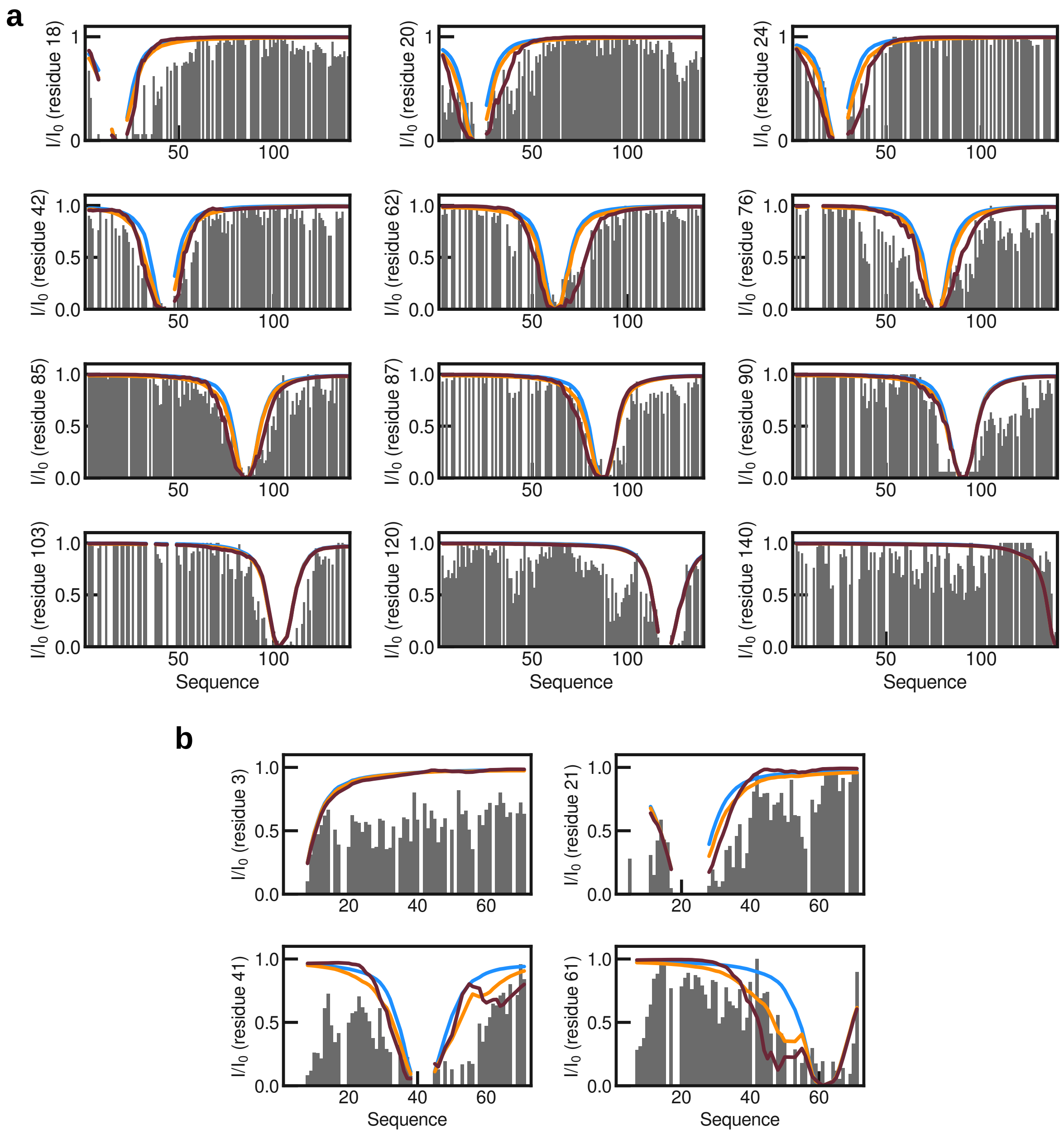


**Supplementary Figure 7: Analysis of Paramagnetic Relaxation Enhancement (PRE) data**. Intensity ratio profiles of PRE experiments on (**a**) $\alpha$-synuclein and (**b**) ACTR. Experimetal data is colored in grey, random coil, bAIes, and bAIes-N in blue, orange, and dark red, respectively.

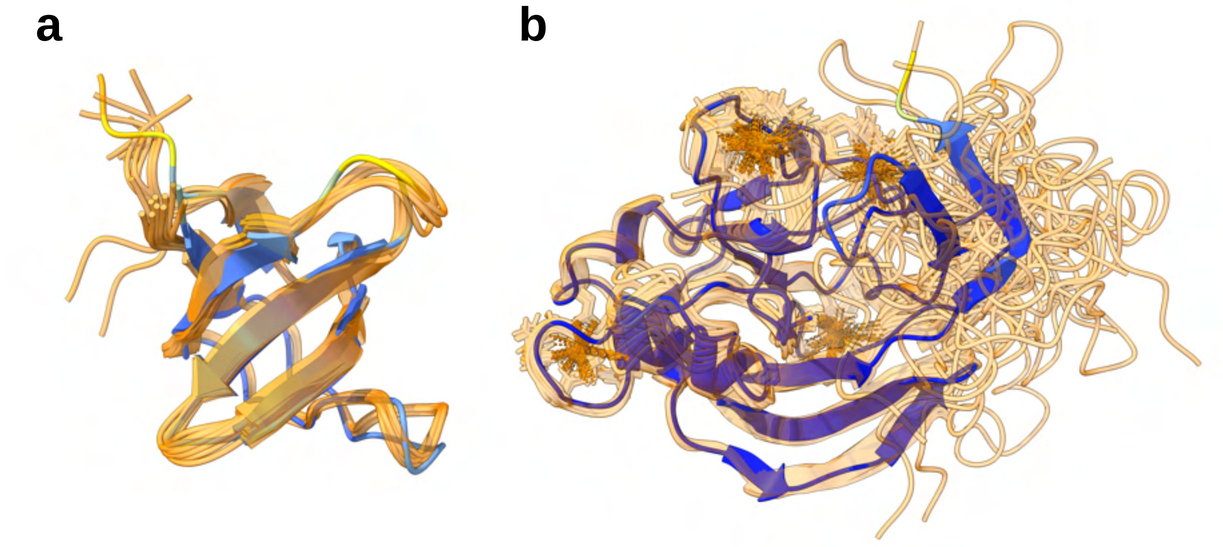


**Supplementary Figure 8: Examples of AF2 prediction errors**. **a**, drkN AF2 model superimposed to the ensemble of structures of the folded state derived by NMR (orange, PDB code 2A36).[3] **b**, spm_FrpC AF2 model superimposed with the ensemble of structures in complex with Calcium ions derived by NMR (orange, PDB code 6SJW).[4]

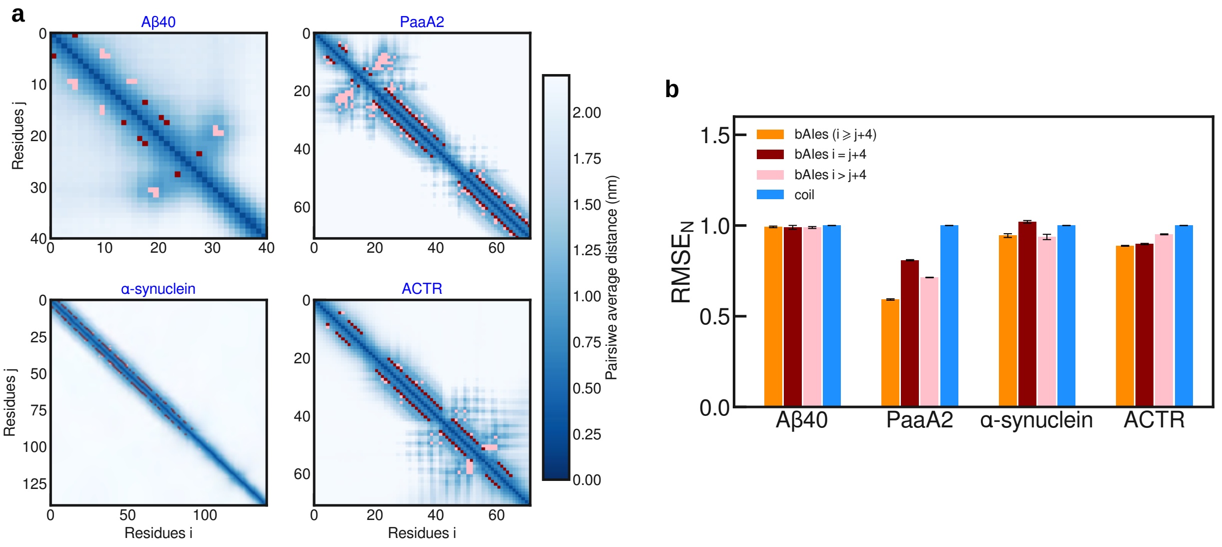


**Supplementary Figure 9: Effect of local and non local predicted contacts on ensemble modeling.** **a**, Distogram average distance matrices for A$\beta$40, PaaA2, ACTR and $\alpha$-synuclein. In dark red are the residue pairs separated by 4 residues, in pink are the residue pairs separated by more than 4 residues. **b**, RMSE$_N$ bar plots of A$\beta$40, PaaA2, ACTR, and $\alpha$-synuclein for the bAIes ensembles obtained with all restraints ($i \geq j+4$, orange), with only $i = j+4$ restraints (dark red), with only $i > j+4$ (pink), and with no restraint (random coil model, blue).

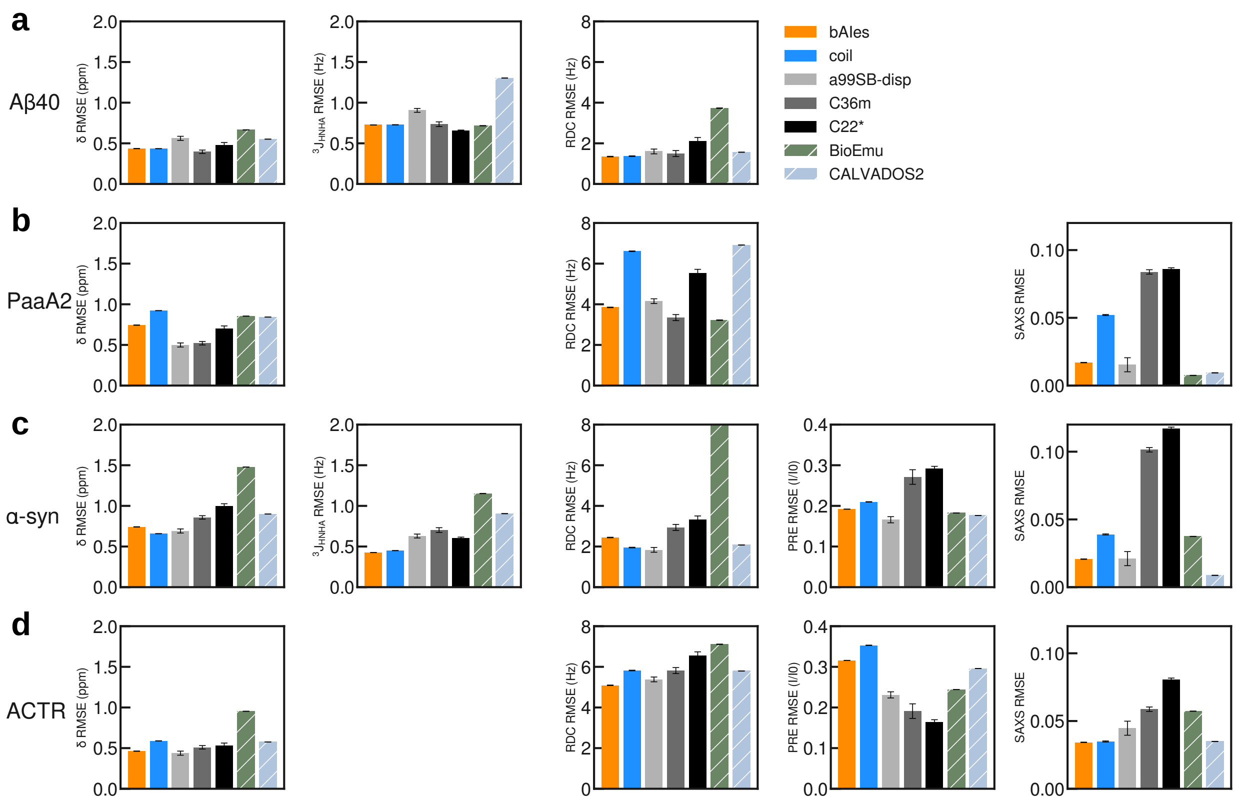


**Supplementary Figure 10: Ensemble fit to each individual experimental data type.** Average RMSEs for each of the available experimental data type (chemical shifts, J-couplings, RDCs, PREs and SAXS) for four IDPs studied in this work: (**a**) A$\beta$40, (**b**) PaaA2 , (**c**) $\alpha$-synuclein , and (**d**) ACTR . This analysis is performed for the random coil and bAIes ensembles as well for ensembles generated with state-of-the-art atomistic MD force fields, CALVADOS2, and BioEmu.

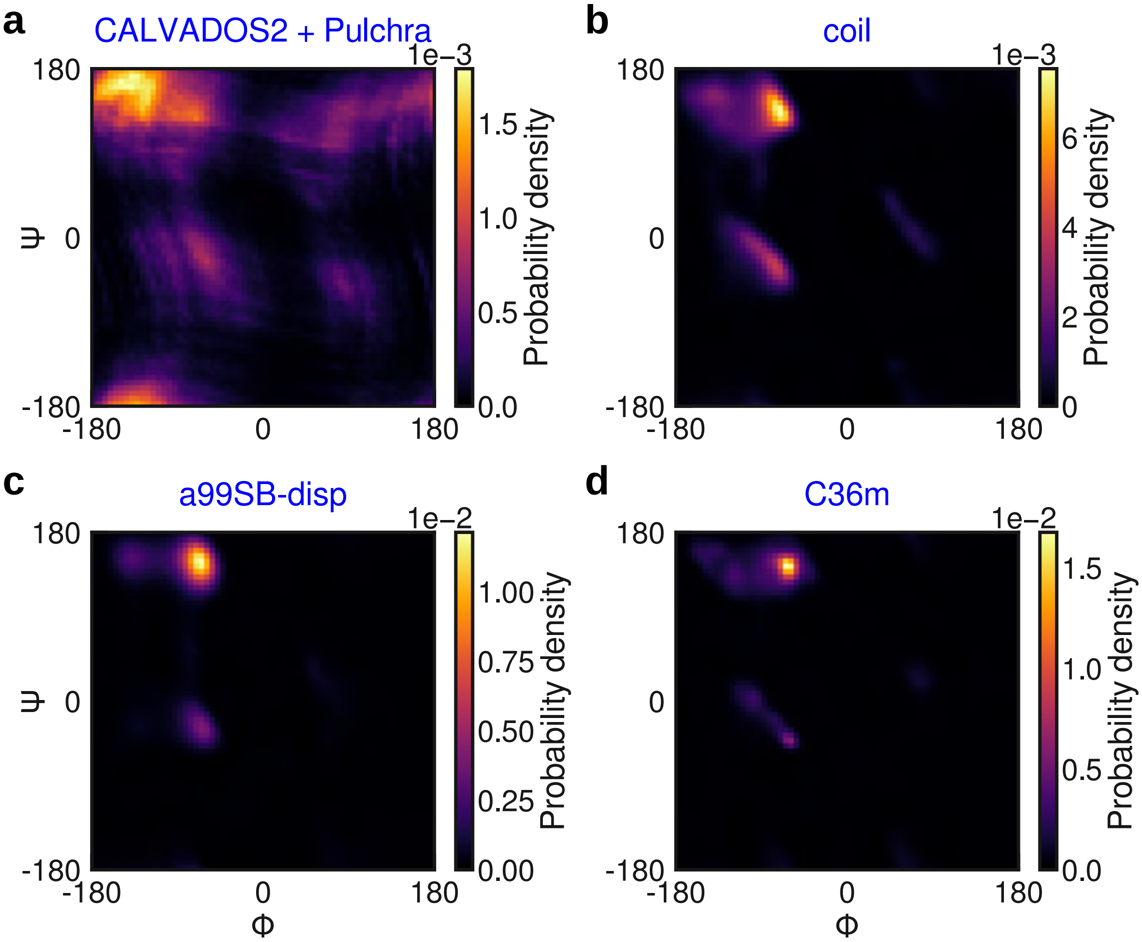


**Supplementary Figure 11: Ramachandran plots with implicit or explicit backbone representations.** Ramachandran plots of $\alpha$-synuclein obtained with (**a**) CALVADOS2,[5] after conversion to all-atom representation with Pulchra,[6] (**b**) our random coil model, (**c**) the amber99SB-disp force field,[2] and (**d**) the CHARMM36m force field.[7]

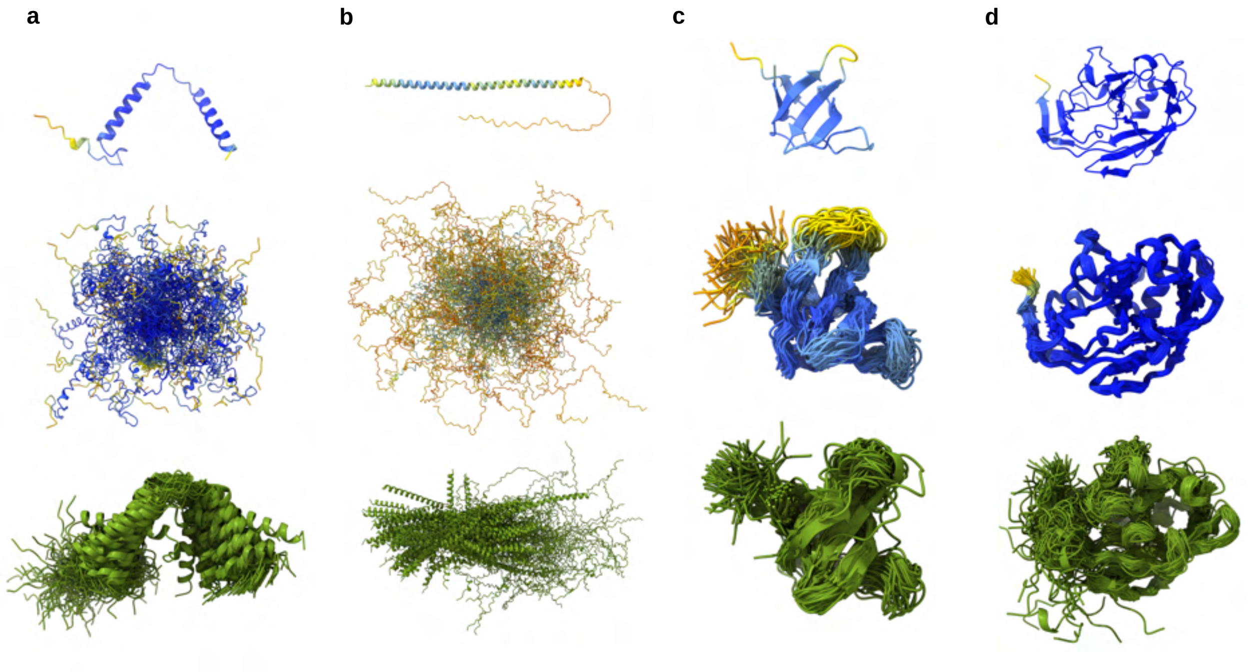


**Supplementary Figure 12: Examples of challenging IDPs for AF2-based structural modeling**. AF2 models (top), bAIes ensembles (middle) and BioEmu ensembles (bottom) for (**a**) PaaA2, (**b**) $\alpha$-synuclein, (**c**) drkN, and (**d**) spm_FrpC.

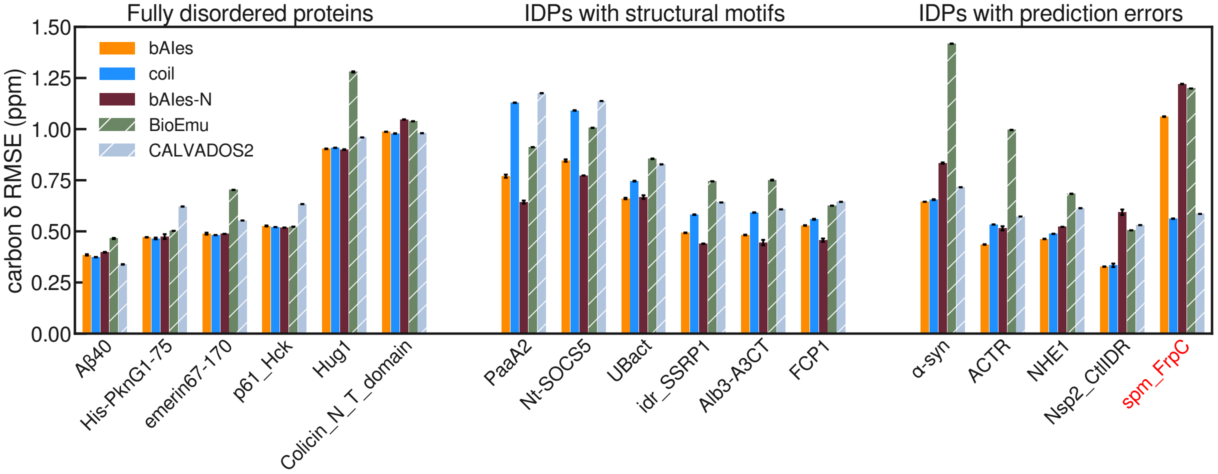


**Supplementary Figure 13: Chemical shifts analysis**. Chemical shift RMSEs averaged over $C_\alpha$, $C_\beta$ and $C'$ for the IDPs studied in this work, calculated on ensembles generated from different IDP models: bAIes, bAIes-N, coil, BioEmu, and CALVADOS2. The IDPs are classified into three groups: fully disordered proteins (left), IDPs with structural motifs (center), and proteins with AF2 prediction errors (left).

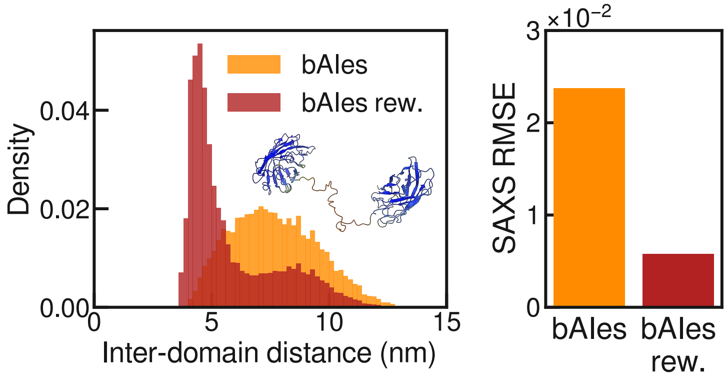


**Supplementary Figure 14: Analysis of GS8 inter-domain distance distributions.** Distribution of the distance between the two centers of mass of the GFPs domains for the bAIes (orange) and reweighted bAIes (red) ensembles of GS8. The center of mass of each GFP domain is defined by residues 1 to 227 and 270 to 486. Right panel: RMSE of the calculated SAXS profiles with respect to the experimental data for the bAIes (orange) and reweighted bAIes (red) ensembles.

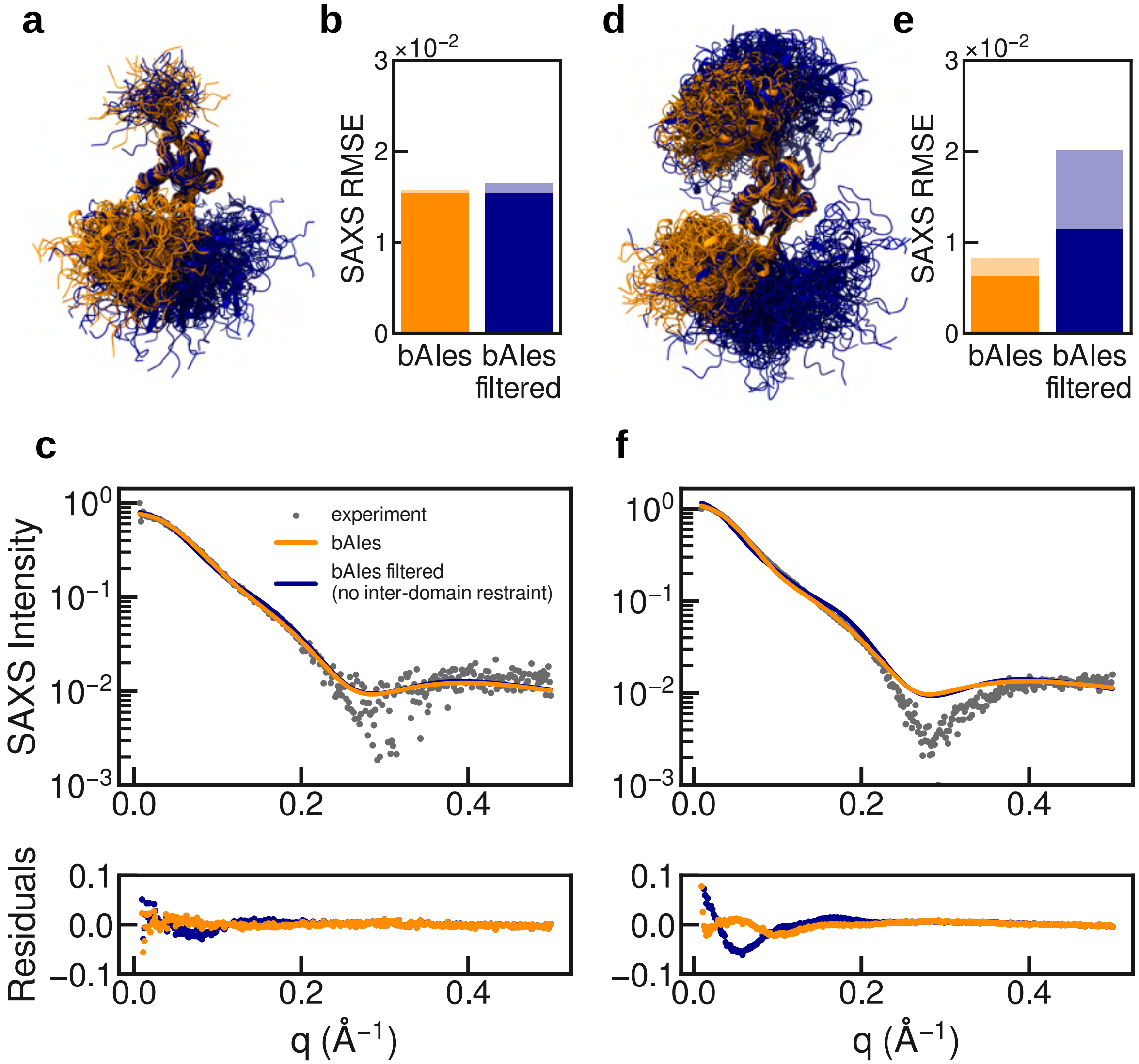


**Supplementary Figure 15: Analysis of contact propensity of polyubiquitins folded domains.** (**a**) and (**d**), bAIes ensembles of Ubq2 and Ubq3, respectively, obtained with (orange) and without (blue) inter-domain contacts predicted by AF2. (**b**) and (**e**), RMSE of the predicted SAXS profile with respect to the experimental data for the bAIes ensembles with (orange) and without (blue) inter-domain contacts. The RMSE bars of the original and reweighted ensembles are colored in light and dark tones, respectively. (**c**) and (**f**), SAXS profiles of Ubq2 and Ubq3, respectively, for the bAIes ensembles with (orange) and without (blue) inter-domain contacts, along with the experimental SAXS profile (grey points). SAXS intensities are shown in logarithmic scale. Bottom panels: difference between experimental and theoretical profiles (residuals).

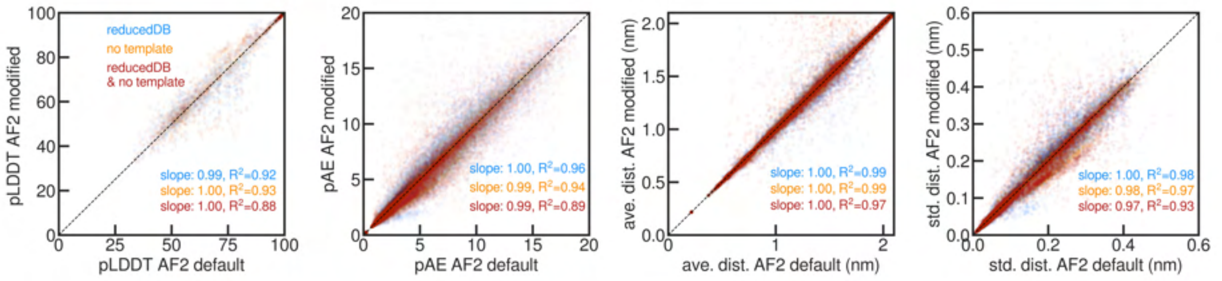


**Supplementary Figure 16: Effect of AF2 parameters on IDPs modeling**. AF2 output pLDDT, pAE, distogram average distance and standard deviation with modified settings as a function of the default AF2 settings. Results obtained with a reduced MSA, with no templates, and with both reduced MSA and no templates are represented in blue, orange and red, respectively. These plot were obtained using the AF2 predictions of all the IDPs studied in this work, except for the multi-domain proteins. In this analysis, only distograms with little information in the last bin (<10%) were kept to minimize inaccuracies in the calculated distance averages and standard deviations.

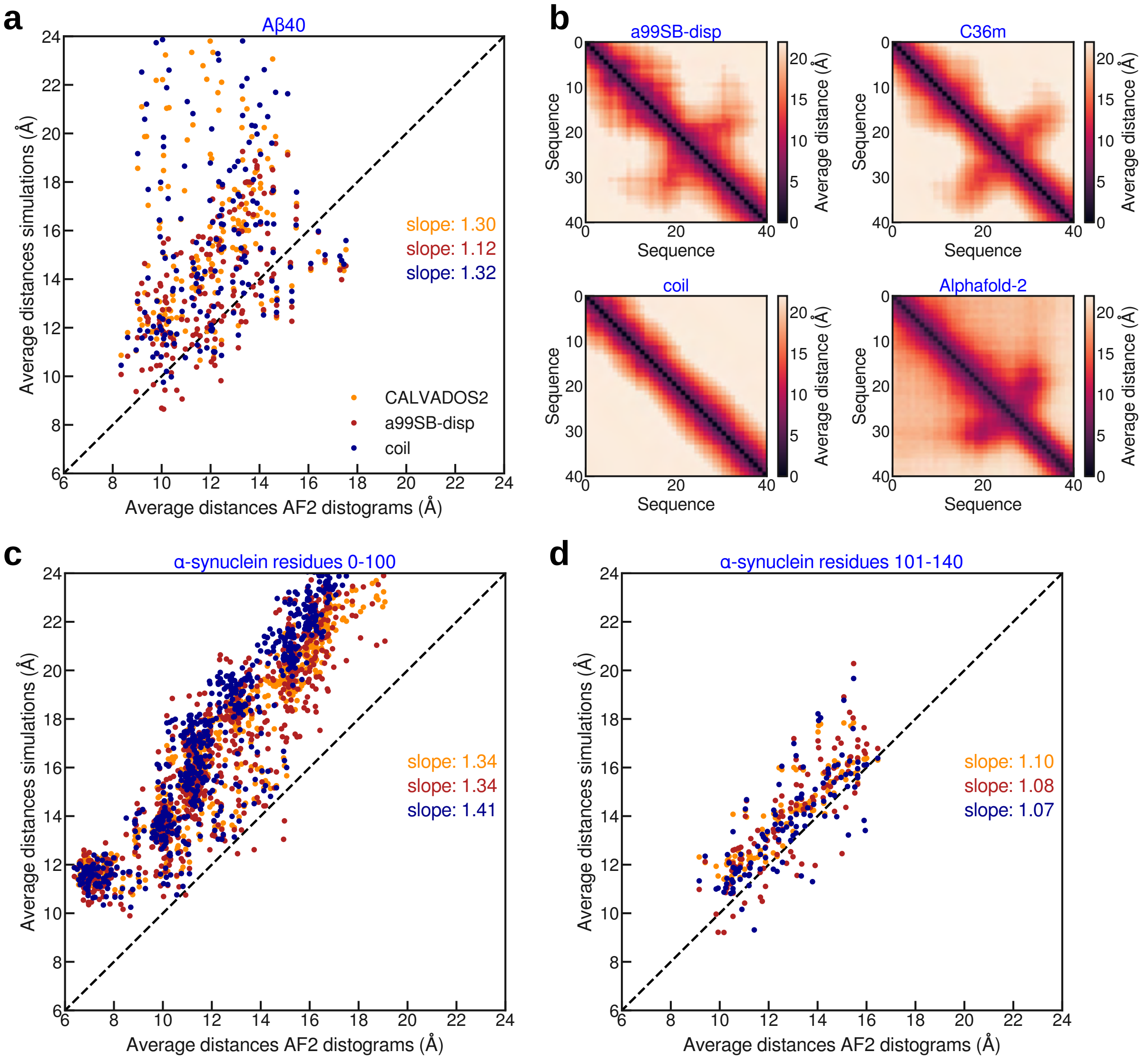


**Supplementary Figure 17: Correlations between average distances from AF2 distograms and molecular simulations.** **a**, Correlation plots of the $C_\beta$-$C_\beta$ average distances of A$\beta$40 computed from random coil (blue), CALVADOS2 (orange), and amber99SB-disp (red) ensembles with respect to the average distances calculated from the AF2 distograms. **b**, Average distance matrices for A$\beta$40 calculated from the amber99SB-disp (top left), CHARMM36m (top right), and random coil (bottom left) ensembles as well as computed from AF2 distograms (bottom right). **c** and **d**, Correlation plots as in panel (**a**) for $\alpha$-synuclein, separately for (**c**) residues 1-100 and (**d**) residues 101-140. For (**a**), (**b**) and (**c**), only the amino acids pairs separated in sequence by at least 4 residues and with AF2 distograms with last bin scarcely populated ($<5\%$) are shown.

### Supplementary Tables

**Supplementary Table 1:** List of simulations performed or analyzed in this work.

| system | SIMULATIONS | coil, bAIes, bAIes-N | C22* C36m a99SB-disp | BioEmu | CALVADOS2 |
|---|---|---|---|---|---|
| **A$\beta$40** | simulation time | 2 $\mu$s | 30 $\mu$s | | |
| | number of conformers | 20000 | 30000 | 25857 | 20000 |
| **PaaA2** | simulation time | 2 $\mu$s | 30 $\mu$s | | |
| | number of conformers | 20000 | 30000 | 25222 | 20000 |
| **$\alpha$-synuclein** | simulation time | 2 $\mu$s | 30 $\mu$s | | |
| | number of conformers | 20000 | 30000 | 18953 | 20000 |
| **ACTR** | simulation time | 2 $\mu$s | 30 $\mu$s | | |
| | number of conformers | 20000 | 30000 | 24264 | 20000 |
| **drkN** | simulation time | 2 $\mu$s | 30 $\mu$s | | |
| | number of conformers | 20000 | 30000 | 29061 | 20000 |
| **emerin67-170** | simulation time | 2 $\mu$s | - | | |
| | number of conformers | 20000 | - | 18546 | 20000 |
| **p61_Hck** | simulation time | 2 $\mu$s | - | | |
| | number of conformers | 20000 | - | 23315 | 20000 |
| **His-PknG1-75** | simulation time | 2 $\mu$s | - | | |
| | number of conformers | 20000 | - | 21618 | 20000 |
| **Colicin_NT_domain** | simulation time | 2 $\mu$s | - | | |
| | number of conformers | 20000 | - | 12912 | 20000 |
| **Hug1** | simulation time | 2 $\mu$s | - | | |
| | number of conformers | 20000 | - | 22902 | 20000 |
| **Nt-SOCS5** | simulation time | 2 $\mu$s | - | | |
| | number of conformers | 20000 | - | 26537 | 20000 |
| **UBact** | simulation time | 2 $\mu$s | - | | |
| | number of conformers | 20000 | - | 19342 | 20000 |
| **idr_SSRP1** | simulation time | 2 $\mu$s | - | | |
| | number of conformers | 20000 | - | 20659 | 20000 |
| **Alb3-A3CT** | simulation time | 2 $\mu$s | - | | |
| | number of conformers | 20000 | - | 12517 | 20000 |
| **FCP1** | simulation time | 2 $\mu$s | - | | |
| | number of conformers | 20000 | - | 20233 | 20000 |
| **NHE1** | simulation time | 2 $\mu$s | - | | |
| | number of conformers | 20000 | - | 19434 | 20000 |
| **Nsp2_CtlIDR** | simulation time | 2 $\mu$s | - | | |
| | number of conformers | 20000 | - | 26940 | 20000 |
| **spm_FrpC** | simulation time | 2 $\mu$s | - | | |
| | number of conformers | 20000 | - | 26356 | 20000 |
| **GS8** | simulation time | 1 $\mu$s | - | - | - |
| | number of conformers | 10000 | - | - | - |
| **Ubq2** | simulation time | 1 $\mu$s | - | - | - |
| | number of conformers | 10000 | - | - | - |
| **Ubq3** | simulation time | 1 $\mu$s | - | - | - |
| | number of conformers | 10000 | - | - | - |

**Supplementary Table 2:** List of experimental data used in this work.

| system | EXPERIMENTAL DATA | CS | RDC | J | PRE | SAXS | Exp. data refs. |
|---|---|---|---|---|---|---|---|
| **A$\beta$40** | Nb. datasets | 4 | 1 | 1 | - | - | $\delta$,[8] $RDC_{NH}$[9] |
| | Nb. data points | 176 | 25 | 36 | - | - | $^3J_{HNHA}$[10] |
| **PaaA2** | Nb. datasets | 6 | 1 | - | - | 1 | all[11] |
| | Nb. data points | 400 | 25 | - | - | 78 | |
| **$\alpha$-synuclein** | Nb. datasets | 5 | 1 | 2 | 12 | 1 | $\delta$,[12] $RDC_{NH}$,[13] $^3J$[14] |
| | Nb. data points | 621 | 101 | 125 | 1220 | 37 | PREs,[15–18] SAXS[19] |
| **ACTR** | Nb. datasets | 5 | 1 | - | 4 | 1 | NMR[20] |
| | Nb. data points | 304 | 57 | - | 203 | 48 | SAXS[21] |
| **emerin67-170** | Nb. datasets | 3 | - | - | - | - | $\delta$[22] |
| | Nb. data points | 297 | - | - | - | - | |
| **p61_Hck** | Nb. datasets | 5 | - | - | - | - | $\delta$[23] |
| | Nb. data points | 360 | - | - | - | - | |
| **His-PknG1-75** | Nb. datasets | 5 | - | - | - | - | $\delta$[24] |
| | Nb. data points | 203 | - | - | - | - | |
| **Colicin_NT_domain** | Nb. datasets | 4 | - | - | - | - | $\delta$[25] |
| | Nb. data points | 322 | - | - | - | - | |
| **Hug1** | Nb. datasets | 5 | - | - | - | - | $\delta$[26] |
| | Nb. data points | 344 | - | - | - | - | |
| **Nt-SOCS5** | Nb. datasets | 5 | - | - | - | - | $\delta$[27] |
| | Nb. data points | 266 | - | - | - | - | |
| **UBact** | Nb. datasets | 5 | - | - | - | - | $\delta$[28] |
| | Nb. data points | 250 | - | - | - | - | |
| **idr_SSRP1** | Nb. datasets | 4 | - | - | - | - | $\delta$[29] |
| | Nb. data points | 303 | - | - | - | - | |
| **Alb3-A3CT** | Nb. datasets | 4 | - | - | - | - | $\delta$[30] |
| | Nb. data points | 262 | - | - | - | - | |
| **FCP1** | Nb. datasets | 5 | - | - | - | - | $\delta$[31] |
| | Nb. data points | 374 | - | - | - | - | |
| **NHE1** | Nb. datasets | 4 | - | - | - | - | $\delta$[32] |
| | Nb. data points | 441 | - | - | - | - | |
| **Nsp2_CtlIDR** | Nb. datasets | 4 | - | - | - | - | $\delta$[33] |
| | Nb. data points | 165 | - | - | - | - | |
| **spm_FrpC** | Nb. datasets | 5 | - | - | - | - | $\delta$[34] |
| | Nb. data points | 759 | - | - | - | - | |
| **GS8** | Nb. datasets | - | - | - | - | 1 | SAXS[35] |
| | Nb. data points | - | - | - | - | 1213 | |
| **Ubq2** | Nb. datasets | - | - | - | - | 1 | SAXS[36] |
| | Nb. data points | - | - | - | - | 345 | |
| **Ubq3** | Nb. datasets | - | - | - | - | 1 | SAXS[36] |
| | Nb. data points | - | - | - | - | 343 | |

**Supplementary Table 3:** Chemical Shift RMSE (*ppm*) for fully disordered IDPs.

| Protein | model | $C_\alpha$ | $C'$ | $C_\beta$ | $N$ | $HN$ | $HA$ |
|---|---|---|---|---|---|---|---|
| | coil | 0.329 | | 0.419 | 1.052 | 0.244 | 0.127 |
| **A$\beta$40** | bAIes | 0.347 | | 0.423 | 1.051 | 0.233 | 0.121 |
| | bAIes-N | 0.343 | | 0.452 | 1.181 | 0.248 | 0.127 |
| | coil | 0.509 | | 0.454 | 0.904 | | |
| **emerin67-140** | bAIes | 0.524 | | 0.454 | 0.931 | | |
| | bAIes-N | 0.517 | | 0.458 | 0.920 | | |
| | coil | 0.416 | 0.681 | 0.466 | 1.115 | | 0.174 |
| **p61_Hck** | bAIes | 0.421 | 0.685 | 0.473 | 1.122 | | 0.175 |
| | bAIes-N | 0.408 | 0.676 | 0.471 | 1.114 | | 0.173 |
| | coil | 0.428 | 0.676 | 0.292 | 1.352 | | 0.104 |
| **His-PknG1-75** | bAIes | 0.431 | 0.681 | 0.301 | 1.360 | | 0.105 |
| | bAIes-N | 0.437 | 0.678 | 0.309 | 1.375 | | 0.111 |
| | coil | 0.452 | 1.924 | 0.559 | 0.718 | | |
| **Colicin_NT_domain** | bAIes | 0.455 | 1.921 | 0.585 | 0.754 | | |
| | bAIes-N | 0.500 | 1.936 | 0.705 | 0.889 | | |
| | coil | 0.583 | 0.600 | 1.543 | 1.238 | | 0.178 |
| **Hug1** | bAIes | 0.579 | 0.593 | 1.539 | 1.204 | | 0.175 |
| | bAIes-N | 0.566 | 0.577 | 1.557 | 1.226 | | 0.171 |

*Note: Areas are empty when there is no available data.*
*The averages are performed over the heavy nuclei.*

**Supplementary Table 4:** Chemical Shift RMSE (*ppm*) for IDPs with residual structure propensities.

| **Protein** | model | $C_\alpha$ | $C'$ | $C_\beta$ | $N$ | $HN$ | $HA$ |
|---|---|---|---|---|---|---|---|
| **PaaA2** | coil | 1.412 | 1.413 | 0.562 | 1.720 | 0.199 | 0.222 |
| | bAIes | 0.859 | 0.933 | 0.518 | 1.732 | 0.247 | 0.172 |
| | bAIes-N | 0.752 | 0.643 | 0.534 | 2.199 | 0.347 | 0.174 |
| **Nt-SOCS5** | coil | 1.334 | | 0.849 | 2.411 | | 0.373 |
| | bAIes | 0.919 | | 0.774 | 1.641 | | 0.329 |
| | bAIes-N | 0.763 | | 0.783 | 1.611 | | 0.248 |
| **UBact** | coil | 0.711 | 0.764 | 0.762 | 1.330 | | 0.218 |
| | bAIes | 0.643 | 0.621 | 0.718 | 1.313 | | 0.194 |
| | bAIes-N | 0.724 | 0.561 | 0.718 | 1.634 | | 0.189 |
| **idr_SSRP1** | coil | 0.653 | 0.724 | 0.368 | 1.263 | | |
| | bAIes | 0.553 | 0.612 | 0.314 | 1.052 | | |
| | bAIes-N | 0.550 | 0.512 | 0.256 | 1.398 | | |
| **Alb3-A3CT** | coil | 0.685 | | 0.499 | 1.139 | | 0.198 |
| | bAIes | 0.540 | | 0.425 | 1.079 | | 0.175 |
| | bAIes-N | 0.476 | | 0.414 | 1.268 | | 0.169 |
| **FCP1** | coil | 0.581 | 0.669 | 0.427 | 1.103 | | 0.135 |
| | bAIes | 0.541 | 0.631 | 0.413 | 1.044 | | 0.126 |
| | bAIes-N | 0.447 | 0.531 | 0.393 | 1.253 | | 0.120 |

*Note: Areas are empty when there is no available data.*
*The averages are performed over the heavy nuclei.*

**Supplementary Table 5:** Chemical Shift RMSE (*ppm*) for IDPs with errors in AF2 predictions.

| Protein | model | $C_\alpha$ | $C'$ | $C_\beta$ | $N$ | $HN$ | $HA$ |
|---|---|---|---|---|---|---|---|
| | coil | 0.367 | 0.496 | 1.101 | 1.171 | 0.152 | |
| **$\alpha$-synuclein** | bAIes | 0.454 | 0.435 | 1.044 | 1.593 | 0.181 | |
| | bAIes-N | 1.021 | 0.475 | 1.006 | 3.302 | 0.330 | |
| | coil | 0.635 | 0.645 | 0.320 | 1.136 | 0.211 | |
| **ACTR** | bAIes | 0.493 | 0.532 | 0.282 | 0.838 | 0.168 | |
| | bAIes-N | 0.617 | 0.485 | 0.443 | 1.968 | 0.200 | |
| | coil | 0.425 | 0.663 | 0.375 | 0.992 | | |
| **NHE1** | bAIes | 0.397 | 0.628 | 0.363 | 1.075 | | |
| | bAIes-N | 0.514 | 0.621 | 0.431 | 1.725 | | |
| | coil | 0.335 | 0.428 | 0.239 | 1.528 | | |
| **Nsp2_CtlIDR** | bAIes | 0.335 | 0.374 | 0.273 | 1.937 | | |
| | bAIes-N | 0.849 | 0.496 | 0.435 | 3.630 | | |
| | coil | 0.609 | 0.740 | 0.337 | 1.059 | | 0.132 |
| **spm_FrpC** | bAIes | 0.978 | 1.011 | 1.194 | 2.372 | | 0.267 |
| | bAIes-N | 1.163 | 1.090 | 1.410 | 2.758 | | 0.334 |

*Note: Areas are empty when there is no available data.*
*The averages are performed over the heavy nuclei.*

**Supplementary Table 6:** RMSE for each experimental data type available, for A$\beta$40 across the different IDPs models tested in this work.

| **data** | **ensemble** | coil | bAIes | bAIes-N | C22* | C36m | a99SB-disp |
|---|---|---|---|---|---|---|---|
| $C_\alpha$ (ppm) | original | 0.329 | 0.347 | 0.343 | 0.444 | 0.403 | 0.472 |
| $C_\alpha$ (ppm) | reweighted | 0.261 | 0.282 | 0.269 | 0.333 | 0.304 | 0.357 |
| $C_\beta$ (ppm) | original | 0.419 | 0.423 | 0.452 | 0.414 | 0.347 | 0.404 |
| $C_\beta$ (ppm) | reweighted | 0.337 | 0.343 | 0.364 | 0.345 | 0.244 | 0.263 |
| $N$ (ppm) | original | 1.052 | 1.051 | 1.181 | 1.149 | 0.914 | 1.543 |
| $N$ (ppm) | reweighted | 0.770 | 0.795 | 0.877 | 0.846 | 0.613 | 1.227 |
| $H_N$ (ppm) | original | 0.244 | 0.233 | 0.248 | 0.292 | 0.225 | 0.295 |
| $H_N$ (ppm) | reweighted | 0.222 | 0.218 | 0.226 | 0.261 | 0.217 | 0.268 |
| $H_\alpha$ (ppm) | original | 0.127 | 0.121 | 0.127 | 0.091 | 0.088 | 0.093 |
| $H_\alpha$ (ppm) | reweighted | 0.105 | 0.101 | 0.104 | 0.075 | 0.035 | 0.063 |
| RDC (Hz) | original | 1.368 | 1.348 | 1.438 | 2.103 | 1.495 | 1.599 |
| RDC (Hz) | reweighted | 0.001 | 0.002 | 0.001 | 0.060 | 1.133 | 0.166 |
| $^3J_{HNHA}$ | original | 0.728 | 0.726 | 0.736 | 0.653 | 0.735 | 0.905 |
| $^3J_{HNHA}$ | reweighted | 0.470 | 0.476 | 0.456 | 0.485 | 0.438 | 0.777 |
| RMSE$_N$ | original | 1.000 | 0.992 | 1.038 | 1.168 | 1.003 | 1.191 |
| RMSE$_N$ | reweighted | 0.549 | 0.557 | 0.559 | 0.590 | 0.743 | 0.766 |

**Supplementary Table 7:** RMSE for each experimental data type available, for PaaA2 across the different atomistic IDP force fields tested in this work.

| **data** | **ensemble** | coil | bAIes | bAIes-N | C22* | C36m | a99SB-disp |
|---|---|---|---|---|---|---|---|
| $C_\alpha$ (ppm) | original | 1.412 | 0.859 | 0.752 | 0.862 | 0.756 | 0.661 |
| $C_\alpha$ (ppm) | reweighted | 1.343 | 0.601 | 0.585 | 0.525 | 0.477 | 0.468 |
| $C$ (ppm) | original | 1.413 | 0.933 | 0.643 | 0.834 | 0.564 | 0.644 |
| $C$ (ppm) | reweighted | 1.337 | 0.693 | 0.485 | 0.523 | 0.377 | 0.458 |
| $C_\beta$ (ppm) | original | 0.562 | 0.518 | 0.534 | 0.548 | 0.490 | 0.422 |
| $C_\beta$ (ppm) | reweighted | 0.489 | 0.413 | 0.444 | 0.431 | 0.403 | 0.355 |
| $N$ (ppm) | original | 1.720 | 1.732 | 2.199 | 1.557 | 1.014 | 0.985 |
| $N$ (ppm) | reweighted | 1.458 | 1.544 | 2.066 | 1.109 | 0.673 | 0.731 |
| $H_N$ (ppm) | original | 0.199 | 0.247 | 0.347 | 0.240 | 0.165 | 0.171 |
| $H_N$ (ppm) | reweighted | 0.177 | 0.209 | 0.331 | 0.183 | 0.090 | 0.116 |
| $H_\alpha$ (ppm) | original | 0.222 | 0.172 | 0.174 | 0.174 | 0.135 | 0.113 |
| $H_\alpha$ (ppm) | reweighted | 0.203 | 0.132 | 0.138 | 0.119 | 0.107 | 0.081 |
| RDC (Hz) | original | 6.608 | 3.846 | 3.122 | 5.536 | 3.344 | 4.153 |
| RDC (Hz) | reweighted | 6.288 | 2.499 | 2.687 | 3.577 | 2.472 | 2.825 |
| SAXS | original | 0.052 | 0.017 | 0.016 | 0.086 | 0.084 | 0.015 |
| SAXS | reweighted | 0.029 | 0.003 | 0.003 | 0.076 | 0.075 | 0.004 |
| $\text{RMSE}_N$ | original | 1.000 | 0.592 | 0.580 | 1.111 | 0.919 | 0.509 |
| $\text{RMSE}_N$ | reweighted | 0.837 | 0.409 | 0.473 | 0.893 | 0.771 | 0.332 |

**Supplementary Table 8:** RMSE for each experimental data type available, for $\alpha$-synuclein across the different atomistic IDP force fields tested in this work.

| **data** | **ensemble** | coil | bAIes | bAIes-N | C22* | C36m | a99SB-disp |
|---|---|---|---|---|---|---|---|
| $C_\alpha$ (ppm) | original | 0.367 | 0.454 | 1.021 | 0.764 | 0.610 | 0.506 |
| $C_\alpha$ (ppm) | reweighted | 0.370 | 0.435 | 0.962 | 0.636 | 0.559 | 0.513 |
| $C$ (ppm) | original | 0.496 | 0.435 | 0.475 | 0.644 | 0.570 | 0.314 |
| $C$ (ppm) | reweighted | 0.464 | 0.407 | 0.424 | 0.568 | 0.485 | 0.237 |
| $C_\beta$ (ppm) | original | 1.101 | 1.044 | 1.006 | 1.267 | 1.275 | 1.043 |
| $C_\beta$ (ppm) | reweighted | 1.041 | 1.000 | 0.962 | 1.189 | 1.156 | 0.981 |
| $N$ (ppm) | original | 1.171 | 1.593 | 3.302 | 2.042 | 1.622 | 1.457 |
| $N$ (ppm) | reweighted | 1.134 | 1.403 | 3.203 | 1.513 | 1.488 | 1.416 |
| $H_N$ (ppm) | original | 0.152 | 0.181 | 0.330 | 0.266 | 0.204 | 0.136 |
| $H_N$ (ppm) | reweighted | 0.148 | 0.163 | 0.313 | 0.215 | 0.147 | 0.137 |
| $^3J_{HNHA}$ (Hz) | original | 0.661 | 0.653 | 1.070 | 0.961 | 1.137 | 1.110 |
| $^3J_{HNHA}$ (Hz) | reweighted | 0.547 | 0.435 | 1.029 | 0.810 | 0.999 | 1.063 |
| $^3J_{CC}$ (Hz) | original | 0.239 | 0.198 | 0.257 | 0.256 | 0.266 | 0.147 |
| $^3J_{CC}$ (Hz) | reweighted | 0.180 | 0.112 | 0.197 | 0.210 | 0.196 | 0.071 |
| RDC (Hz) | original | 1.949 | 2.444 | 7.090 | 3.337 | 2.952 | 1.835 |
| RDC (Hz) | reweighted | 1.377 | 1.387 | 6.942 | 2.878 | 2.604 | 1.441 |
| $PRE_{18}$ $(I/I_0)$ | original | 0.250 | 0.229 | 0.238 | 0.339 | 0.359 | 0.216 |
| $PRE_{18}$ $(I/I_0)$ | reweighted | 0.229 | 0.197 | 0.229 | 0.313 | 0.267 | 0.198 |
| $PRE_{20}$ $(I/I_0)$ | original | 0.256 | 0.230 | 0.208 | 0.276 | 0.357 | 0.228 |
| $PRE_{20}$ $(I/I_0)$ | reweighted | 0.235 | 0.205 | 0.198 | 0.268 | 0.250 | 0.209 |
| $PRE_{24}$ $(I/I_0)$ | original | 0.116 | 0.095 | 0.102 | 0.265 | 0.294 | 0.104 |
| $PRE_{24}$ $(I/I_0)$ | reweighted | 0.093 | 0.066 | 0.096 | 0.256 | 0.240 | 0.086 |
| $PRE_{42}$ $(I/I_0)$ | original | 0.182 | 0.136 | 0.125 | 0.405 | 0.302 | 0.131 |
| $PRE_{42}$ $(I/I_0)$ | reweighted | 0.160 | 0.108 | 0.106 | 0.334 | 0.260 | 0.113 |
| $PRE_{62}$ $(I/I_0)$ | original | 0.155 | 0.135 | 0.131 | 0.236 | 0.306 | 0.104 |
| $PRE_{62}$ $(I/I_0)$ | reweighted | 0.136 | 0.104 | 0.127 | 0.179 | 0.225 | 0.088 |
| $PRE_{76}$ $(I/I_0)$ | original | 0.301 | 0.273 | 0.229 | 0.416 | 0.244 | 0.190 |
| $PRE_{76}$ $(I/I_0)$ | reweighted | 0.278 | 0.242 | 0.186 | 0.362 | 0.230 | 0.187 |
| $PRE_{85}$ $(I/I_0)$ | original | 0.177 | 0.161 | 0.132 | 0.318 | 0.235 | 0.159 |
| $PRE_{85}$ $(I/I_0)$ | reweighted | 0.152 | 0.122 | 0.102 | 0.281 | 0.203 | 0.129 |
| $PRE_{87}$ $(I/I_0)$ | original | 0.160 | 0.147 | 0.157 | 0.259 | 0.251 | 0.133 |
| $PRE_{87}$ $(I/I_0)$ | reweighted | 0.130 | 0.105 | 0.106 | 0.229 | 0.180 | 0.096 |
| $PRE_{90}$ $(I/I_0)$ | original | 0.252 | 0.238 | 0.237 | 0.276 | 0.324 | 0.175 |
| $PRE_{90}$ $(I/I_0)$ | reweighted | 0.226 | 0.193 | 0.184 | 0.256 | 0.230 | 0.160 |
| $PRE_{103}$ $(I/I_0)$ | original | 0.185 | 0.181 | 0.183 | 0.173 | 0.163 | 0.151 |
| $PRE_{103}$ $(I/I_0)$ | reweighted | 0.161 | 0.143 | 0.134 | 0.196 | 0.151 | 0.131 |
| $PRE_{120}$ $(I/I_0)$ | original | 0.259 | 0.254 | 0.255 | 0.325 | 0.256 | 0.206 |
| $PRE_{120}$ $(I/I_0)$ | reweighted | 0.247 | 0.237 | 0.220 | 0.280 | 0.242 | 0.209 |
| $PRE_{140}$ $(I/I_0)$ | original | 0.226 | 0.224 | 0.224 | 0.195 | 0.192 | 0.194 |
| $PRE_{140}$ $(I/I_0)$ | reweighted | 0.219 | 0.210 | 0.207 | 0.181 | 0.190 | 0.203 |
| SAXS | original | 0.039 | 0.021 | 0.021 | 0.117 | 0.101 | 0.021 |
| SAXS | reweighted | 0.022 | 0.010 | 0.008 | 0.109 | 0.090 | 0.013 |
| $\text{RMSE}_N$ | original | 1.000 | 0.944 | 1.668 | 1.805 | 1.653 | 0.889 |
| $\text{RMSE}_N$ | reweighted | 0.853 | 0.714 | 1.595 | 1.695 | 1.516 | 0.807 |

**Supplementary Table 9:** RMSE for each experimental data type available, for ACTR across the different atomistic IDP force fields tested in this work.

| **data** | **ensemble** | coil | bAIes | bAIes-N | C22* | C36m | a99SB-disp |
|---|---|---|---|---|---|---|---|
| $C_\alpha$ (ppm) | original | 0.635 | 0.493 | 0.617 | 0.558 | 0.499 | 0.469 |
| $C_\alpha$ (ppm) | reweighted | 0.577 | 0.412 | 0.521 | 0.294 | 0.329 | 0.350 |
| $C$ (ppm) | original | 0.645 | 0.532 | 0.485 | 0.436 | 0.485 | 0.413 |
| $C$ (ppm) | reweighted | 0.533 | 0.457 | 0.411 | 0.325 | 0.225 | 0.332 |
| $C_\beta$ (ppm) | original | 0.320 | 0.282 | 0.443 | 0.327 | 0.293 | 0.285 |
| $C_\beta$ (ppm) | reweighted | 0.250 | 0.195 | 0.379 | 0.216 | 0.185 | 0.230 |
| $N$ (ppm) | original | 1.136 | 0.838 | 1.968 | 1.166 | 1.065 | 0.843 |
| $N$ (ppm) | reweighted | 0.930 | 0.616 | 1.753 | 0.640 | 0.776 | 0.616 |
| $H_N$ (ppm) | original | 0.211 | 0.168 | 0.200 | 0.165 | 0.207 | 0.179 |
| $H_N$ (ppm) | reweighted | 0.190 | 0.143 | 0.175 | 0.115 | 0.163 | 0.151 |
| RDC (Hz) | original | 5.820 | 5.091 | 6.977 | 6.555 | 5.815 | 5.381 |
| RDC (Hz) | reweighted | 4.762 | 2.547 | 6.669 | 4.442 | 4.330 | 4.078 |
| $PRE_3$ $(I/I_0)$ | original | 0.372 | 0.368 | 0.363 | 0.119 | 0.124 | 0.191 |
| $PRE_3$ $(I/I_0)$ | reweighted | 0.356 | 0.362 | 0.363 | 0.096 | 0.105 | 0.172 |
| $PRE_{21}$ $(I/I_0)$ | original | 0.328 | 0.290 | 0.290 | 0.179 | 0.228 | 0.211 |
| $PRE_{21}$ $(I/I_0)$ | reweighted | 0.296 | 0.265 | 0.281 | 0.150 | 0.174 | 0.204 |
| $PRE_{31}$ $(I/I_0)$ | original | 0.421 | 0.362 | 0.392 | 0.178 | 0.247 | 0.295 |
| $PRE_{31}$ $(I/I_0)$ | reweighted | 0.391 | 0.353 | 0.391 | 0.128 | 0.169 | 0.286 |
| $PRE_{61}$ $(I/I_0)$ | original | 0.290 | 0.243 | 0.290 | 0.180 | 0.165 | 0.227 |
| $PRE_{61}$ $(I/I_0)$ | reweighted | 0.279 | 0.245 | 0.291 | 0.151 | 0.160 | 0.227 |
| SAXS | original | 0.035 | 0.034 | 0.038 | 0.081 | 0.059 | 0.045 |
| SAXS | reweighted | 0.034 | 0.034 | 0.036 | 0.072 | 0.061 | 0.040 |
| RMSE$_N$ | original | 1.000 | 0.888 | 1.095 | 1.197 | 1.025 | 0.909 |
| RMSE$_N$ | reweighted | 0.948 | 0.770 | 1.096 | 0.974 | 0.910 | 0.815 |

(8) Hou, L.; Shao, H.; Zhang, Y.; Li, H.; Menon, N. K.; Neuhaus, E. B.; Brewer, J. M.; Byeon, I.-J. L.; Ray, D. G.; Vitek, M. P.; Iwashita, T.; Makula, R. A.; Przybyla, A. B.; Zagorski, M. G. Solution NMR Studies of the A(140) and A(142) Peptides Establish

that the Met35 Oxidation State Affects the Mechanism of Amyloid Formation. *Journal of the American Chemical Society* **2004**, *126*, 1992–2005.

(15) Dedmon, M. M.; Lindorff-Larsen, K.; Christodoulou, J.; Vendruscolo, M.; Dobson, C. M. Mapping Long-Range Interactions in -Synuclein using Spin-Label NMR and

Ensemble Molecular Dynamics Simulations. *Journal of the American Chemical Society* **2005**, *127*, 476–477.

(22) Samson, C.; Herrada, I.; Celli, F.; Theillet, F.-X.; Zinn-Justin, S. 1H, 13C and 15N

backbone resonance assignment of the intrinsically disordered region of the nuclear envelope protein emerin. *Biomolecular NMR assignments* **2016**, *10*, 179–182.

(29) Hashimoto, M.; Kodera, N.; Tsunaka, Y.; Oda, M.; Tanimoto, M.; Ando, T.; Morikawa, K.; Tate, S.-i. Phosphorylation-coupled intramolecular dynamics of unstruc-

tured regions in chromatin remodeler FACT. *Biophysical Journal* **2013**, *104*, 2222–2234.
